# Targeting calpastatin pharmacologically restores synaptic proteolysis and preserves motor neurons survival and function in C9orf72 ALS

**DOI:** 10.64898/2025.12.08.692907

**Authors:** Léa Lescouzères, Zoé Butti, Mathilde Chaineau, Daniel Young, Zhipeng You, Michael Nicouleau, Carol X-Q Chen, Ghazal Haghi, Nathalia Aprahamian, Charlotte Zaouter, Antoine Dufour, Thomas M. Durcan, Shunmoogum A. Patten

## Abstract

A hexanucleotide repeat expansion (GGGGCC) in the *C9orf72* gene is the most prevalent genetic cause of ALS, with early neuromuscular junction (NMJ) dysfunction being a key pathological feature. Current therapies provide only limited symptomatic relief, underscoring the need for targeted, mechanism-based interventions.

Using a C9orf72 ALS zebrafish model (C9-miR) and patient-derived induced pluripotent stem cell (iPSC) motor neurons, we identified significant downregulation of calpastatin, the endogenous inhibitor of calpains, a calcium-dependent protease family implicated in neurodegeneration.

We demonstrate that restoring calpastatin activity with a cell-permeable calpastatin-derived peptide or the small molecule, calpeptin, ameliorates locomotor deficits and NMJ dysfunction in the C9-miR zebrafish model. These interventions enhance synaptic vesicle turnover and quantal release at the NMJ while improving motor neuron excitability and synaptic integrity in iPSC-derived motor neurons.

N-terminomic/TAILS mass spectrometry revealed direct calpain-mediated cleavage of synaptic proteins in motor neurons derived from C9orf72 patients. Proteolysis of novel ALS-relevant synaptic and axonal proteins is prevented by calpeptin and calpastatin peptide treatments.

Our findings establish the calpastatin as a pivotal regulator of synaptic function in C9orf72-associated ALS and identify it as a promising therapeutic target, offering a novel strategy to restore synaptic transmission and potentially halt disease progression.

## INTRODUCTION

Amyotrophic lateral sclerosis (ALS) is a neurodegenerative disease that causes motoneuron loss, leading to progressive paralysis and death, typically within ∼5years of diagnosis ^1^. More than two dozen genes have been linked to ALS ^2–4^, with a hexanucleotide repeat expansion (GGGGCC) *C9orf72* being the most common genetic cause (accounting for ∼40% familial and 8-10% of sporadic cases) ^5,6^. Current ALS treatments provide only marginal clinical benefits, underscoring the urgent need for effective disease-modifying therapies.

Disruption of calcium homeostasis and synaptic dysfunction are central features of ALS physiopathology ^7,8^. Calpastatin, the endogenous inhibitor of calcium-activated cysteine proteases, plays a critical role in maintaining proteolytic balance in neurons. It contains four inhibitory domains, enabling simultaneous inhibition of multiple calpain protease molecules ^9,10^. Importantly, calpastatin depletion has been associated with neuronal vulnerability in neurodegenerative conditions ^11–15^, suggesting that loss of this protective brake contributes to disease progression. In ALS patients and experimental models, altered calcium homeostasis correlate with reduced calpastatin levels and impaired neuronal function ^16,8^.

In the hSOD1^G93A^ ALS mouse model, decreased calpastatin levels in the spinal cord is accompanied by motoneuron degeneration, whereas calpastatin overexpression prevented motoneuron death and promoted their survival, even at later stages of the disease ^14^. Similarly, in slow channel congenital myasthenic syndrome (SCS), excessive calpain activity impairs synaptic transmission at the neuromuscular junction (NMJ) ^17^. Expression of human calpastatin in a mouse model of SCS restored calpain activity, preserved NMJ structure and miniature endplate current, and improved synaptic transmission at NMJ ^17^. Because defects in NMJs are among the earliest pathological features in ALS, strategies aimed at stabilizing and preserving NMJ function may hold therapeutic potential to slow or halt disease progression. Notably, as demonstrated by the ongoing Phase 1 LUMINA trial assessing AMX0114, an antisense oligonucleotide intended to modulate a major driver of axonal degeneration in ALS, which has demonstrated favorable early safety and tolerability in patients, therapeutic approaches targeting proteolytic imbalance are gaining clinical traction. However, the potential of calpastatin in protecting NMJ function in ALS, particularly C9orf72-associated form, remains unexplored.

We recently developed a zebrafish model of C9orf72-related ALS ^18^. Zebrafish is an ideal model for studying ALS due to the high conservation of spinal cord physiology and genetics across vertebrates ^19,20^. The zebrafish C9orf72 ALS model faithfully recapitulates key features of the human disease, including muscle atrophy, motor deficits, TDP-43 pathology, synaptic/NMJ impairments ^18^ and DNA damage ^21^. Notably, we found that the zebrafish *c9orf72* model exhibits a marked decrease in the expression of calpastatin (cast) ^18^, suggesting an early impairment of endogenous proteostatic mechanisms.

In this study, we demonstrate that calpastatin expression is reduced in both the c9orf72 zebrafish model and motor neurons (MNs) derived from newly generated induced pluripotent stem cell (iPSC) lines from ALS patients. We show that pharmacological restoration of calpastatin function, using either the small molecule calpeptin or an active calpastatin peptide, leads to significant improvements in both models. Specifically, treatment ameliorated motor behavior, restored NMJ structure, enhanced unitary synaptic transmission, and normalized synaptic vesicle cycling at NMJ in the c9orf72 zebrafish model. In iPSC-derived MNs, these treatments not only improved survival but also restored synaptic integrity and transmission. Importantly, using an unbiased proteomics and N-terminomics approach, we identify novel ALS-relevant synaptic and axonal proteins and proteolytic substrates directly in disease-relevant motor neurons. This strategy provides a comprehensive and previously unexplored view of calpastatin-dependent dysregulated proteostasis in ALS, and uncover new candidate effectors downstream of calpastatin deficiency in a pathological context. Together, these findings highlight the therapeutic potential of targeting the calpastatin to alleviate NMJ dysfunction in C9orf72-associated ALS.

## RESULTS

### Calpastatin expression is reduced in C9-miR zebrafish and C9orf72 ALS iPSC-derived motor neurons

We previously showed that a zebrafish c9orf72 loss-of-function model recapitulates key ALS phenotypes, including motor deficits, muscle atrophy and TDP-43 pathology ^18^. Proteomic analysis in that study revealed a significant reduction in the expression of calpastatin in the C9orf72 zebrafish ALS model (referred hereafter as C9-miR) (**Fig 1A**).

**Figure 1:**
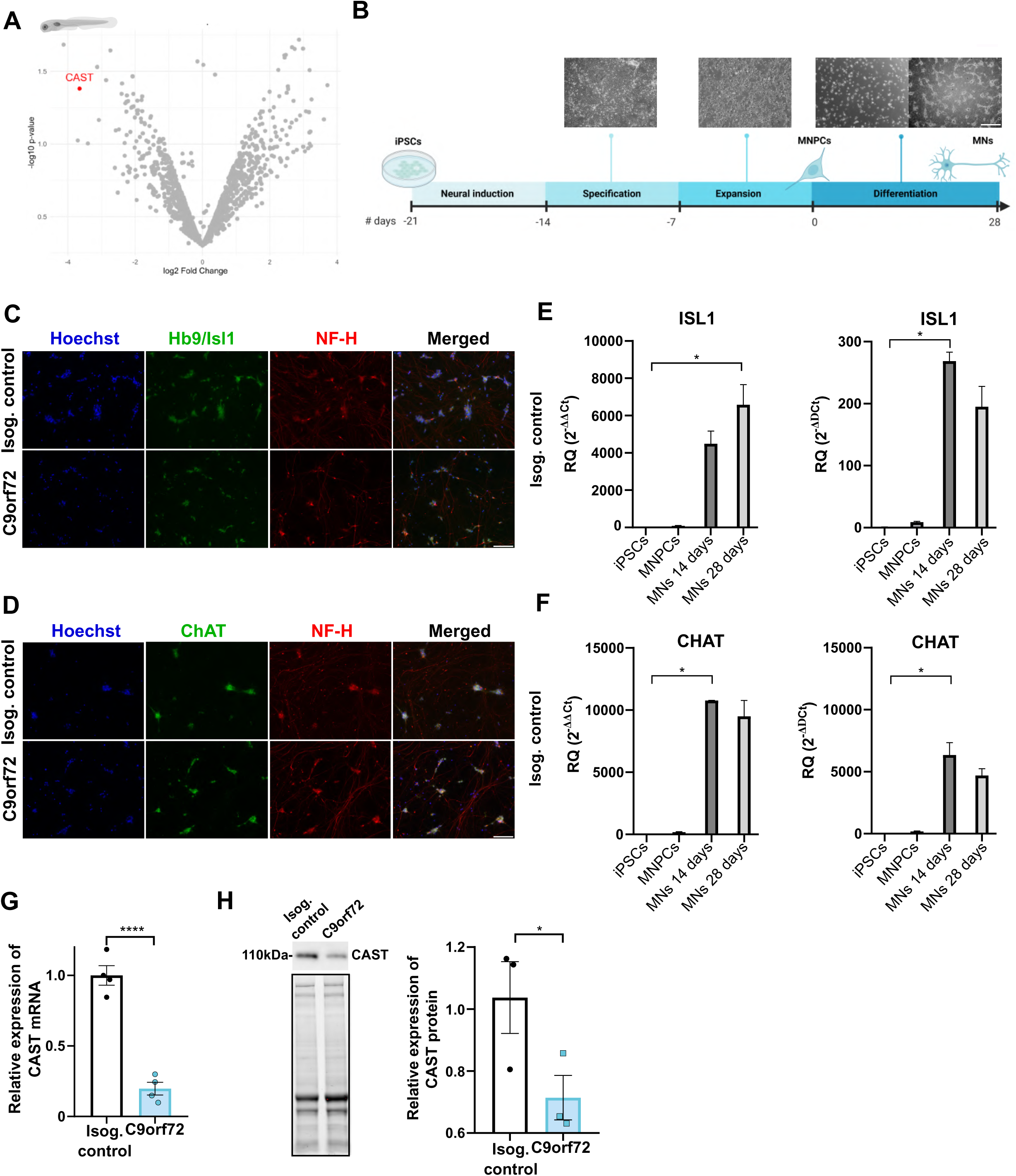
A newly characterized C9orf72 motor neuron model uncovers calpastatin downregulation **(A)** Volcano plot illustrating log₂ fold change versus-log₁₀ p-value for differential expression between control and C9-miR fish (N = 4 per group). Each point represents an individual protein. The downregulation of CAST is highlighted in red. (**B**) Schematic representation of the protocol for sequential differentiation of iPSCs into neuroepithelial progenitors (NEPs), MNPCs, and MNs with representative phase-contrast images of cells along differentiation. Scale bar, 250 μm. (**C,D**) Representative images of C9orf72 isogenic and C9orf72 iPSC-derived MNs visualized by immunochemistry with the combined HB9 and Islet1 MN markers (HB9/ISL1) (**C**) and the choline acetyltransferase protein (ChAT) (**D**), and the neuronal marker neurofilament heavy chain (NF-H), after 28 days of final differentiation. Nuclei were counterstained with Hoechst. Scale bar: 100μm. (**E,F**) Normalized expression levels of ISL1 (**E**), and CHAT (**F**) in iPSCs, MNPCs, and iPSC-derived MNs differentiated for 14 and 28 days from C9Orf72 isogenic (left panel) and C9Orf72 (right panel) lines. Data normalized to Actβ-GAPDH expression. Bar graphs show the mean ± SD. (**G**) Bar graph showing the relative expression of the endogenous Calpastatin gene. mRNA was normalized to GAPDH mRNA level (Kruskal-Wallis test). (**H**) Immunoblot and associated quantification of Calpastatin protein levels. Extractions were performed in MNs harvested after 3 weeks post-plating. Data are represented as mean ± SEM. *P < 0.05, ****P<0.0001.

Given that dysregulated calpastatin-calpain signaling has been implicated in several neurodegenerative diseases, yet its role in C9orf72-associated ALS remains unexplored, we next investigated whether calpastatin dysregulation also occurs in human C9orf72 ALS and contributes to disease pathophysiology. To this end, we examined calpastatin expression in spinal motor neurons (MNs) differentiated form C9orf72 patient-derived IPSC lines and isogenic CRISPR-corrected controls (**Fig Supp. 1, Fig Supp. 2, see materials and methods for details**). MN differentiation was performed using a well-established protocol that recapitulates the sequential developmental progression of spinal MNs ^22,23^ (**Fig 1B**). Briefly, iPSCs were first induced into neuroepithelial progenitors (NEPs), which were next specified into motor neuron progenitor cells (MNPCs) over 6 days. MNPCs were subsequently replated for terminal differentiation into mature MNs. MNs were cultured for 14-and 28-days post-plating to capture two complementary maturation stages, representing early post-mitotic MNs with initial axon extension, and mature MNs, respectively. Differentiation efficiency was confirmed by immunofluorescence and qPCR (**Fig Supp. 3**). At the MNPC stage, we confirmed expression of the neural precursor markers Nestin and Ki67, the latter being an endogenous marker of active cell cycling, thereby confirming the proliferative capacity of cells (**Fig Supp. 3A**). Characterization of the newly generated cell lines was validated by the expected downregulation of the pluripotency markers NANOG and POU5F1 at the MNPC and MN stages compared to the iPSC stage (**Fig Supp. 3B**). In parallel, the motor neuron progenitor OLIG2 marker was upregulated at the MNPC stage relative to both iPSC and MN stages. Finally, expression of the mature MN markers HB9 (MNX1), ISL1, (**Fig 1C,D**) and CHAT was strongly upregulated in mature MNs (**Fig 1E,F**), while being nearly absent in the iPSCs and MNPCs (**Fig 1C-F**). We assessed calpastatin levels and distribution in MNs at 28 days post-plating, a state at which they exhibit more mature electrophysiological and synaptic properties ^24^. Our results showed a significant reduction in calpastatin expression in C9orf72 IPSC-derived MNs compared to isogenic controls, at both the mRNA (**Fig 1G**) and protein (**Fig 1H**) levels.

Calpastatin normally resides near the nuclear invaginations. Upon elevation of intracellular Ca^2+^ concentration, calpastatin undergoes an intracellular redistribution to the cytosol, where it inhibits calpain activity ^25^. Changes in calpastatin’s cellular localization are a well-established marker of its activation and subsequent inhibition of calpain activity ^25,26^. To gain further insight into calpastatin function in C9orf72 ALS, we examined its cellular localization in C9orf72 IPSC-derived MNs and their isogenic controls. Calpastatin is localized predominantly to axons, strongly colocalizing with neurofilament heavy chain (NF-H) in both genotypes (Manders’ coefficient ≈ 1; **Fig Supp. 4A, B**). Interestingly, calpastatin showed markedly stronger nuclear expression in C9orf72 MNs compared to isogenic controls (**Fig Supp. 4A**), accompanied by a significant increase in the nuclear-to-cytoplasmic signal ratio (**Fig Supp. 4C**). Altogether, these data suggest an altered cellular distribution of calpastatin and, consequently, likely dysregulation of calpastatin function in C9orf72 ALS.

### Calpastatin active peptide and the small molecule calpeptin ameliorate motor behavioral phenotype and restore NMJ integrity in C9-miR zebrafish

To determine whether restoring calpastatin function could counteract pathogenic alterations in C9orf72 ALS, we assessed the neuroprotective effects of two complementary approaches in the zebrafish C9-miR ALS model: restoration of calpastatin activity using a cell-permeable calpastatin active peptide (CAST) and pharmacological mimicking the endogenous calpastatin function in calpain activity with the drug calpeptin (Calp) (**Fig 2A**). At 6 dpf, C9-miR larvae exhibit a robust and quantifiable reduction in motor activity, a phenotype previously validated in several zebrafish ALS models ^27–30^. Using this motor impairment as a sensitive readout, we assessed the therapeutic efficacy of CAST and Calp in C9-miR fish (**Fig 2A**). Treatment with both CAST and Calp significantly improved the motor behavior of C9-miR zebrafish (**Fig 2B**), with a marked increase in the total distance travelled, comparable to the levels observed in control fish.

**Figure 2:**
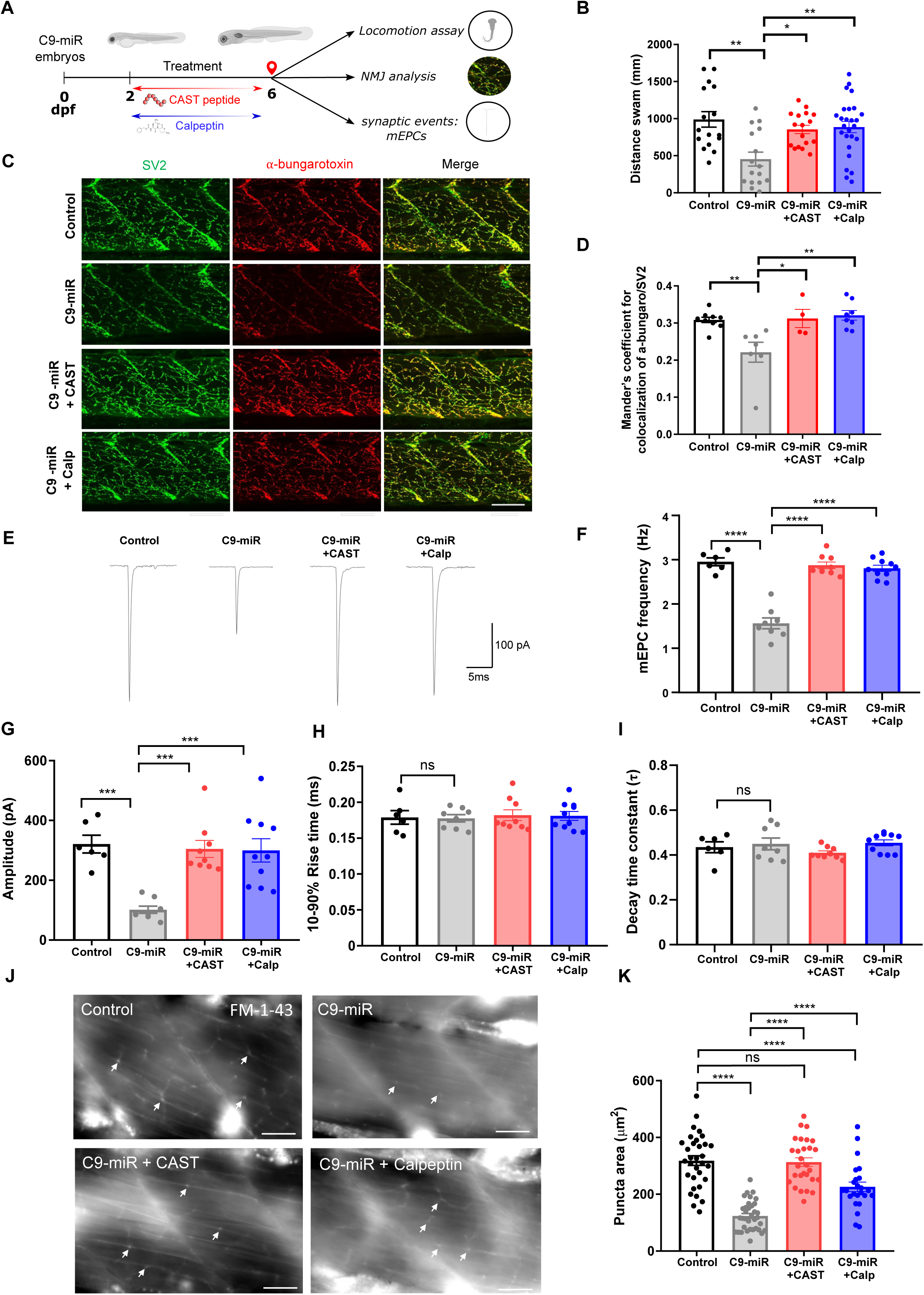
CAST and Calp treatments restore NMJ function and synaptic vesicle dynamics in C9-miR zebrafish. (**A**) Schematic representation of the treatments on C9-miR fish, exposed to CAST peptide and Calpeptin for four days starting at 2 days post fertilization (dpf). All phenotypic results were obtained at 6dpf. (**B**) Treated C9-miR fish displayed improved swimming behavior (CAST: N=3, n=17; Calp: N=3, n=26, 1-way ANOVA). Data are represented as mean ± SEM. (**C**) Representative images of co-immunostaining of zebrafish neuromuscular junction with presynaptic (SV2; green) and postsynaptic (α-bungarotoxin; red) markers in 6 dpf zebrafish showing a rescue of the number of NMJ clusters with both treatments. Scale bar = 100µm. (**D**) Quantification of the colocalizing pre-and post-synaptic markers showing a significant increase of the NMJ number in both treated fish compared to C9-miR (CAST: n=4; Calp: n=8, 1-way ANOVA \**P* < 0.05, \*\**P* < 0.01 (**E**) Representative mEPCs. Fish treated with both compounds show a rescued frequency (**F**) and amplitude (**G**). Rise time (**H**) and decay time (**I**) were not found to be significantly different. (One way ANOVA). (**J**) Representative images of the loading of FM1-43 dye at the NMJ in 6 dpf fish showing a significant rescue of the puncta area indicating a rescue of the synaptic vesicles turnover in treated fish. (**K**) Quantification of FM1-43 puncta area in control (n=32), C9-miR (n=36), C9-mir+Calp (n=24) and C9-miR+CAST (n=28), One way ANOVA. Data are represented as mean ± SEM. ns, not statistically significant; *P < 0.05, **P < 0.01, ***P < 0.001, ****P<0.0001.

We next examined whether CAST and Calp treatments were effective in preserving NMJ structure integrity in C9-miR zebrafish. Immunostaining of NMJ structure in 6 dpf zebrafish C9-miR and control larvae with the pre-synaptic (SV2a) and post-synaptic (α-bungarotoxin) markers revealed structural abnormalities in C9-miR fish compared to controls (**Fig 2C,D**), as we previously reported ^18^. CAST and Calp treatments had a protective effect on NMJ structure in C9-miR larvae (**Fig 2C, D**). These findings imply that calpastatin compensation by Calp and CAST could preserve NMJ structural integrity and motor function in the C9-miR zebrafish.

### CAST and calpeptin improve synaptic transmission and synaptic vesicle release and recycling at NMJs in C9-miR zebrafish

To further characterize the neuroprotective potential of CAST and Calp in our C9-miR zebrafish model, we next assessed whether the observed improvements in NMJ integrity extended to synaptic transmission at the NMJ. To this end, we recorded and analyzed quantal events, focusing on spontaneous miniature end plate currents (mEPCs) at the NMJs (**Fig 2E**). Analysis of mEPCs revealed a significant decrease in both frequency and amplitude (**Fig 2 F, G**) in C9-miR larvae compared to controls. Treatment with CAST or Calp significantly increased mEPCs frequency and amplitude (**Fig 2 F, G**) in C9-miR. We observed no differences in rise time (**Fig 2H**) or decay time constant (**Fig 2I**) kinetics among C9-miR larvae, controls and treated groups. Altogether, our findings indicate that pharmacological restoration of calpastatin function effectively mitigates NMJ functional deficits in a C9orf72 zebrafish ALS model.

We previously reported downregulation of the synaptic vesicle SV2A in C9-miR zebrafish, associated with reduced synaptic vesicle release and recycling ^18^. SV2A is a critical component of the presynaptic release machinery and its level can be restored by calapstatin in models of synaptic dysfunction ^17^. Based on these findings, we sought to further investigate synaptic activity at the NMJ in C9-miR treated with CAST or Calp by measuring synaptic vesicle cycling using the fluorescent styryl dye FM1-43 (**Fig 2J**) ^31,32^. Following depolarization-induced uptake, FM1-43 labeling revealed a marked reduction on fluorescence intensity and puncta size in C9-miR larvae compared to control, indicating a slower exocytotic activity and synaptic vesicle cycle in C9-miR fish (**Fig 2 J, K**). Notably, treatment with Calp or CAST significantly improved FM1-43 uptake in presynaptic terminals of C9-miR larvae compared to untreated C9-miR fish, reflecting improved synaptic vesicle cycling (**Fig 2 J,K**). These findings suggest that calpastatin modulates presynaptic vesicle release and recycling at the NMJ and may ameliorate synaptic defects associated with C9orf72 pathology.

### Calpastatin pharmacological compensation enhances cell survival and restores synaptic integrity and transmission in C9orf72 motor neurons

We next evaluate the therapeutic potential of modulating the calpastatin to improve the survival of MNs derived from C9orf72 ALS patients (**Fig 3A**). Neuronal vulnerability was experimentally induced in MN cultures by neurotrophic factor (NTF) withdrawal, a standard protocol widely used in the ALS research to reveal disease-associated survival deficits and shown to trigger degeneration in ALS MNs carrying C9orf72 ^33,34^ or TDP-43 ^35^ mutations. Viability of MNs differentiated for 4 weeks was assessed using an ATP-based luminescent viability assay. Under basal conditions, C9orf72 MN cultures exhibited survival rates comparable to those of their isogenic controls (**Fig 3B**). However, NTF deprivation reduced C9orf72 MN survival by approximately 20% at 4 weeks post-plating, with no significant effect on controls (**Fig 3B**).

**Figure 3:**
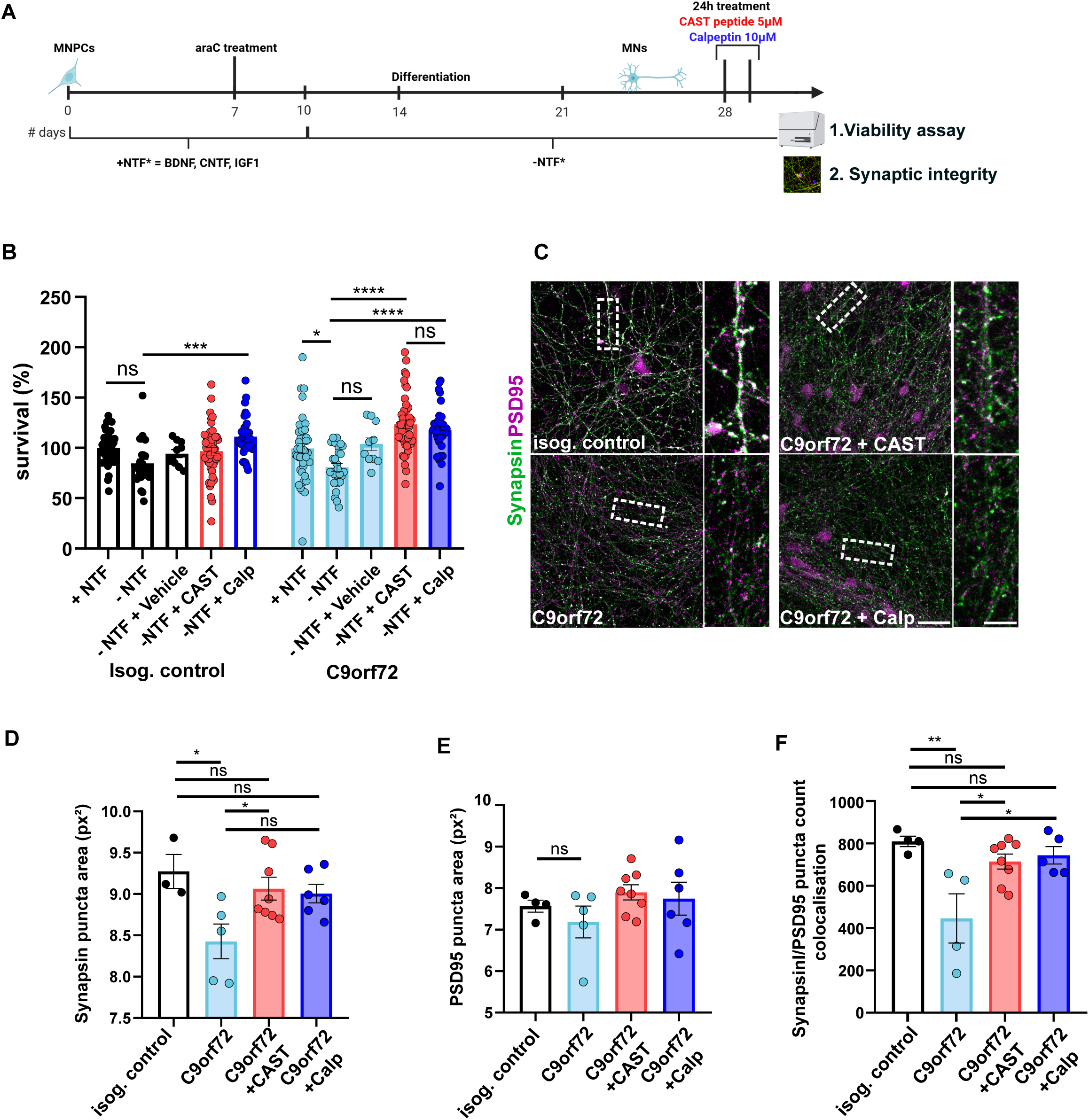
Calpastatin compensation with CAST and Calp treatment improves survival and rescue synaptic architecture in C9orf72 MN cultures. (**A**) Schematic representation of the protocol for MNs treatment with CAST and Calp before viability assay. (**B**) Bar graphs of viability of MN cultures differentiated with and without NTFs at 4 weeks post-plating and after treatment with 5µM active CAST peptide or 10µM calpeptin for 24 hours. (1-way ANOVA). (**C**) Representative images of 4 weeks post-plating MN neurites subjected to immunocytochemistry for synapsin I and PSD95. Scale bar: 10µm. Quantification of the synapsin and PSD95 puncta area (**D-E**) and colocalized puncta count (**F**). Data are represented as mean ± SEM. ns, not statistically significant; *P < 0.05, **P < 0.01, ***P < 0.001, ****P<0.0001.

Treatment with either an active CAST peptide (CAST, 5µM) or calpeptin (Calp, 10µM) for 24 hours significantly improved survival of C9orf72 MNs. Specifically, survival rates reached 117.6% in CAST-treated cultures (C9orf72 + CAST) and 123.5% in Calp-treated cultures (C9orf72 + Calp), compared to approximately 80% in untreated C9orf72 MNs (**Fig 3B**).

Given the restoration of NMJ integrity and synaptic function in vivo, we next assessed synaptic integrity in human C9orf72 MNs. Co-immunostaining for the pre-synaptic marker synapsin I and the post-synaptic marker PSD95 revealed a significant reduction in synapsin+ puncta size in C9orf72 MNs, consistent with a presynaptic defect, while PSD95+ puncta were unchanged (**Fig 3C-E**). We next assessed whether Calp and CAST treatment could preserve synaptic structural integrity. CAST treatment fully rescued synapsin⁺ puncta size, whereas Calp treatment produced a partial rescue (**Fig 3C, D**). Both treatments significantly increased the number of colocalized synapsin⁺/PSD95⁺ puncta, reflecting recovery of overall synaptic organization (**Fig 3F**). Altogether, these findings demonstrate that enhancing calpastatin function preferentially rescues presynaptic defects in C9orf72 MNs.

To determine whether these structural improvements translated into functional recovery, we performed multielectrode array (MEA) ^35^ recordings over a four-week differentiation period (**Fig 4A**). MEA recordings allow for the measurement of extracellular electrical activity in neuronal networks, providing quantitative insights into neuronal excitability and network dynamics. Three key parameters were analyzed: spikes, defined as individual action potentials detected by electrodes; bursts, representing synchronized clusters of spikes indicative of coordinated network activity, and firing rate, reflecting the frequency of spikes per unit time. Notably, at one-week post-plating, C9orf72 mutant cells exhibited hyperexcitability, characterized by a significant increase in the number of spikes per burst and mean firing rate (**Fig 4 B, D-F**). However, this initial hyperactivity was followed by a progressive and significant decline in mean firing rate, number of spikes per burst and burst frequency in the C9orf72 line (**Fig 4B-F**). The hypoactivity ultimately culminated in a complete loss of burst activity, with most electrodes remaining entirely silent in the C9orf72 cultures (**Fig 4C, D-F**).

**Figure 4:**
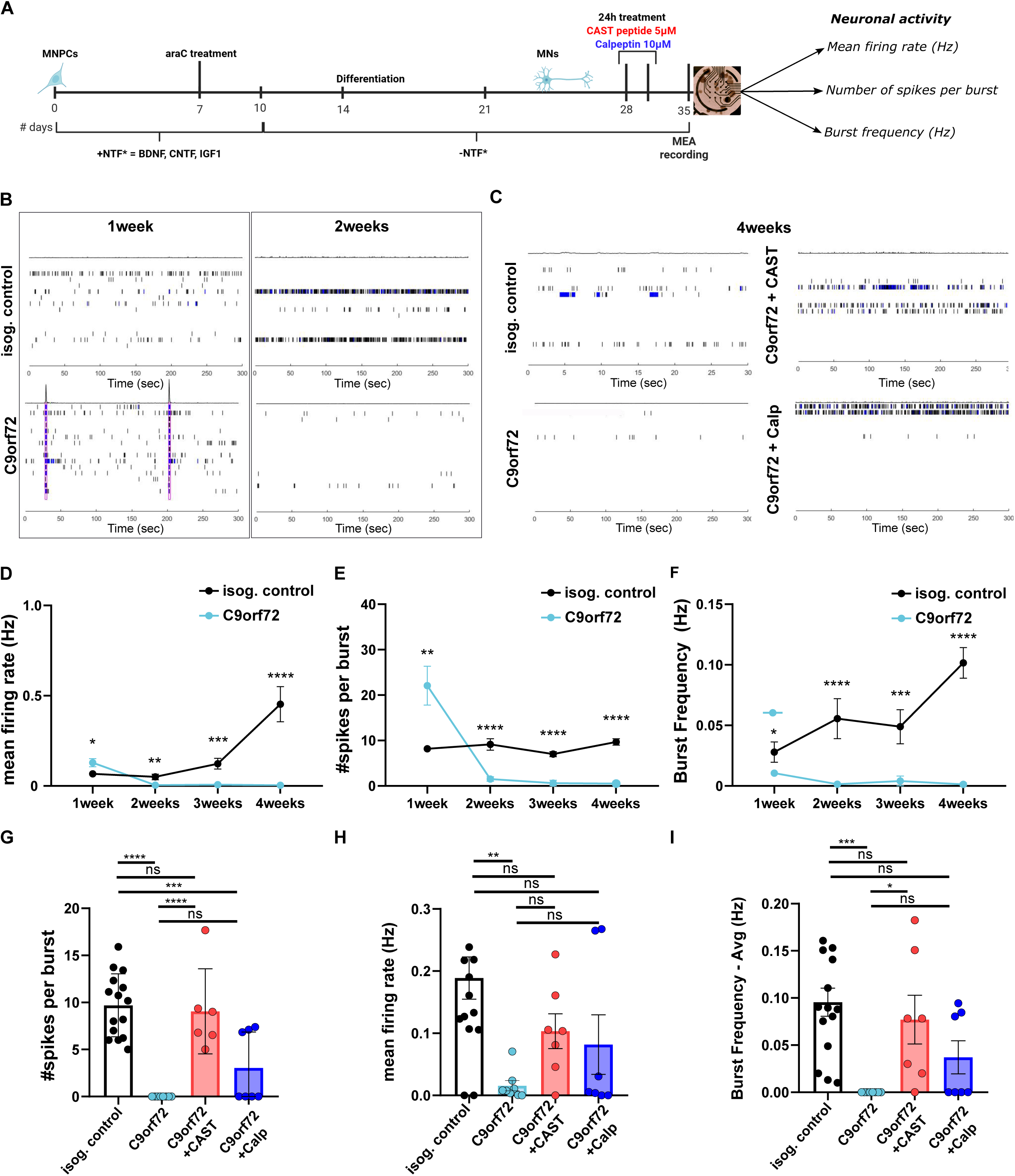
CAST and Calp treatments rescue synaptic architecture and synaptic function in C9orf72 MN cultures. (**A**) Overview of the neuronal activity assay with a muti-electrode array (MEA). Spontaneous neuronal activity of MN cultures differentiated for 1 and 2 weeks (**B**) and 4 weeks with or without treatment (**C**) recorded for 300 s shown as raster plots and spike histograms. Individual spikes are shown in black and bursts are shown in blue. Longitudinal changes in mean firing rate (**D**), number of spikes by burst (**E**) and Burst Frequency average (**F**) of MNs recorded weekly over a span of 4 weeks post-plating. NB: in the absence of burst, values have been artificially adjusted to zero. Quantification de mean firing rate (**G**), number of spikes by burst (**H**) and Burst Frequency average (**I**) of 4 weeks post-plating MNs recorded 1 week after a 24hours-treatment (One-way-ANOVA). Data are represented as mean ± SEM. ns, not statistically significant; *P < 0.05, **P < 0.01, ***P < 0.001, ****P<0.0001.

A single 24-hour treatment with CAST or Calp led to sustained functional improvements. Seven days post-treatment, CAST significantly restored bursting activity, including spikes per burst and burst frequency, whereas untreated C9orf72 MNs remained silent (**Fig 4G-I**). Both compounds also increased the mean firing rate (**Fig 4H**), although these effects did not reach statistical significance. Calpeptin treatment, however, showed more variable effects, consistent with a partial rescue (**Fig 4G-I**). Overall, these findings strongly suggest that, in addition to promoting cell survival, CAST treatments significantly restore spontaneous neuronal activity in C9orf72 motor neurons. The delayed yet sustained recovery following transient treatments suggests long-lasting modulation of neuronal function, potentially through stabilisation of synaptic components.

### Calpastatin expression is also reduced in sporadic ALS iPSC-derived motor neurons

We next examined whether our findings on C9orf72 ALS iPSC-derived MNs could be extended to sporadic ALS. We observed reduced calpastatin expression in MNs derived from a sporadic ALS iPSC line (**Fig Supp 5A, B**). We also found that calpastatin expression was markedly increased in the nucleus of sporadic MNs compared to controls (**Fig Supp 5C**). Treatment with CAST or Calp significantly improved survival of sporadic ALS MNs (**Fig Supp 5D**). These findings demonstrate that enhancing calpastatin activity, either directly via CAST peptide or indirectly through calpain inhibition with calpeptin, effectively mitigates motor neuron death and prolongs viability in sporadic ALS.

Sporadic MNs exhibited a significant decline in mean firing rate, number of spikes per burst and burst frequency as of 1-week post-plating (**Fig Supp 5E-G**). However, neither CAST nor Calp restored the activity of sporadic (**Fig Supp 5H-J**), suggesting that distinct molecular mechanisms underlie synaptic dysfunction in sporadic versus C9orf72 ALS.

### Shotgun proteomics reveals extensive proteome remodeling in C9orf72 MNs and its normalization by CAST and Calp treatment

To comprehensively characterize proteome alterations associated with C9orf72 pathology and assess their modulation by pharmacological treatment, we performed a quantitative shotgun proteomics analysis workflow coupled to LC-MS/MS (**Fig 5A**). Motor neurons derived from isogenic control, C9orf72, C9orf72 + CAST, and C9orf72 + Calp iPSC lines were lysed and isotopically labeled using light formaldehyde, digested with trypsin, and analyzed by high-resolution mass spectrometry.

**Figure 5:**
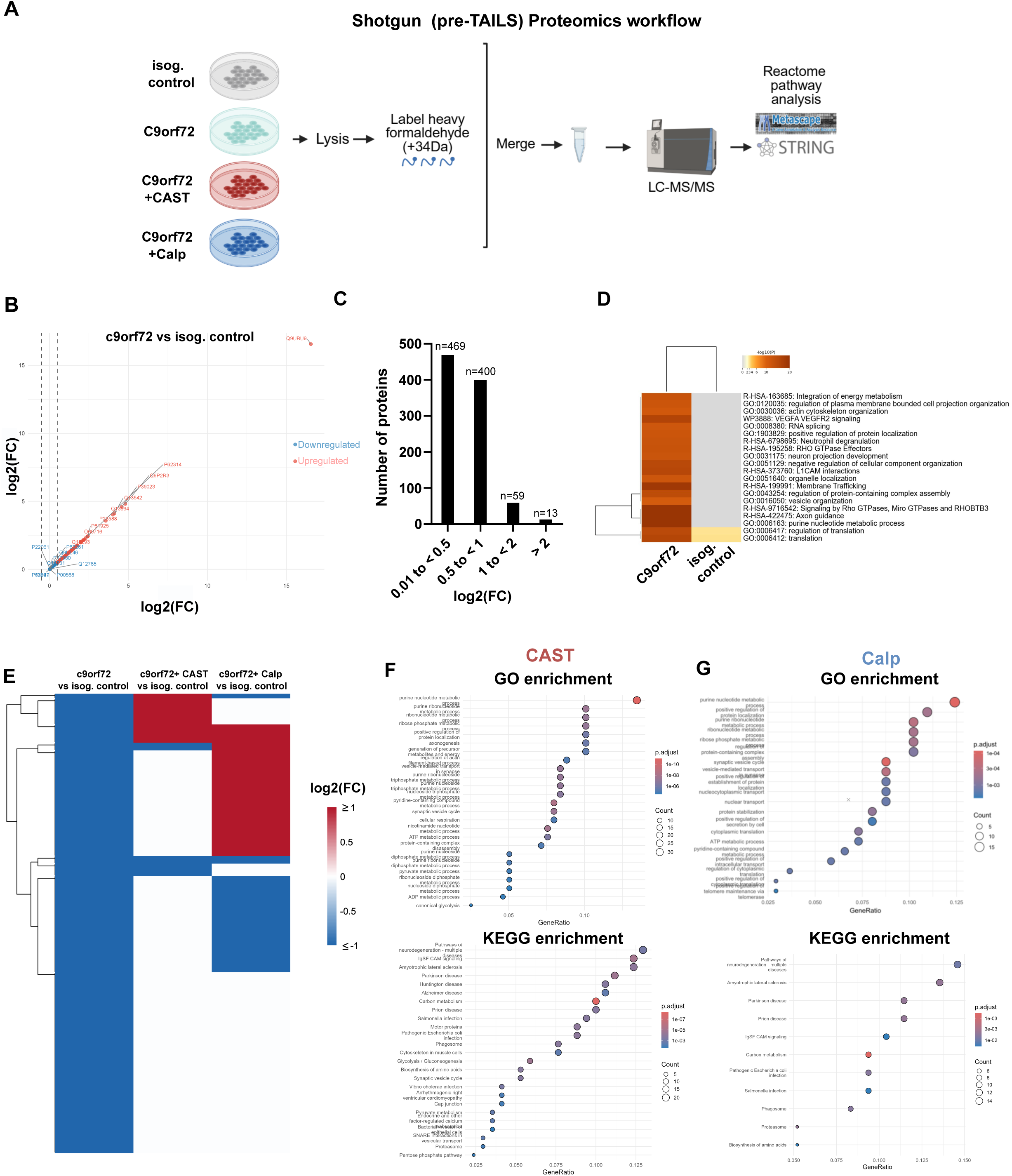
Shotgun proteomic analysis of C9orf72-associated protein alterations and treatment-induced rescue. **(A)** Schematic workflow of shotgun (pre-TAILS) proteomics experimental design. (**B**) Volcano plot showing differentially expressed proteins in C9orf72 compared to isogenic control, with significantly upregulated (red) and downregulated (blue) proteins based on log2 fold change. (**C**) Distribution of log2 fold change values across all proteins identified in the shotgun proteomic dataset. (**D**) Metascape (Zhou et al., 2019) enrichment heatmap showing significantly overrepresented biological terms between C9orf72 and isogenic control motor neurons. (**E**) Heatmap illustrating the shift in protein expression profiles in response to treatment, with proteins downregulated in C9orf72 (blue) and progressively returning toward non-significant log2 fold change values (white) following CAST peptide (middle) and calpeptin (right) treatment, while upregulated proteins are shown in red. (**F-G**) Gene ratio-based GO (top panel) and KEGG (bottom panel) pathway enrichment analyses of proteins whose expression is restored toward non-significant levels after CAST (F) or calpeptin (G) treatment.

Comparative analysis between C9orf72 and isogenic control MNs revealed widespread proteomic alterations, with 469 proteins downregulated and 13 upregulated proteins (meeting a log2 fold-change cutoff) (**Fig 5B, C**). To better understand the molecular processes affected in C9orf72 MNs, we performed pathway enrichment analysis using Metascape. This analysis revealed significant disruption of key cellular pathways, including energy metabolism, ribosomal function and translation, RNA splicing, membrane trafficking, cytoskeletal organization, and axon guidance (**Fig 5D**).

To evaluate potential treatment-induced proteomic rescue, we conducted two comparative analyses: (1) untreated C9orf72 MNs versus C9orf72 MNs treated with CAST, and (2) untreated C9orf72 MNs versus C9orf72 MNs treated with Calp. CAST treatment robustly restored a large subset of dysregulated proteins toward control levels, as evidenced by a pronounced shift of log2FC values toward zero (**Fig. 5E**, middle panel). In constrast, Calp treatment produced a more moderate rescue effect, with fewer proteins returning to baseline expression levels **(Fig. 5E**, right panel).

Functional enrichment of proteins that were downregulated in C9orf72 MNs but restored to near control levels following CAST treatment revealed significant enrichment in key biological processes and pathways, including ribosomal function, energy metabolism, proteasome activity, glycolysis, synaptic vesicle cycling, and neurodegenerative disease pathways, most notably ALS (**Fig 5F**). A highly similar enrichment profile was observed for proteins rescued by Calp treatment (**Fig 5G**), indicationg that both compounds converge on on overlapping mechanisms to restore proteostasis in C9orf72 MNs.

Collectively, these findings demonstrate extensive proteostatic dysfunction in C9orf72 motor neurons. Importantly, they show that CAST and Calp treatments can rescue these disrupted protein networks, an effect consistent with the observed improved neuronal function and survival of C9orf72 MNs when treated with CAST or Calp.

### N-terminomic profiling identifies CAST-and calpeptin-modulated cleavage landscapes in C9orf72 motor neurons

Depletion of calpastatin in C9orf72 MNs can lead to increased protein calpain-dependent proteolysis, promotin pathological protein cleavage and subsequently impair t motor neuron survival and function ^36,37^. To identify proteolytic cleavage events in untreated and treated C9orf72 MNs, we performed N-terminomics using the HYTANE strategy ^38–42^. This approach enables unbiased profiling of both native N-termini arising from translation and “neo” N-termini generated by proteolytic cleavage. Neo-N-termini, which represent cleavage sites, are selectively enriched by removing N-terminals formed after trypsin cleavage using HYTANE method. By enriching and sequencing these N-terminal peptides via mass spectrometry, the strategy provides a direct read-out of protease-mediated processing events and offers insights into how N-terminal modifications affect protein stability, localization, and function.

Proteins from isogenic control MNs, untreated C9orf72 MNs and C9orf72 MNs treated for 24h with either CAST or Calp were labeled with light formaldehyde (+28 Da demethylation) (**Fig 6A**). To identify cleavage sites (neo-N-termini), untreated and treated C9orf72 MNs were analyzed using the N-terminomics/HYTANE protocol, where the N-termini are enriched using undecanal to attach a hydrophobic tag ^43^ (**Fig 6A**).

**Figure 6:**
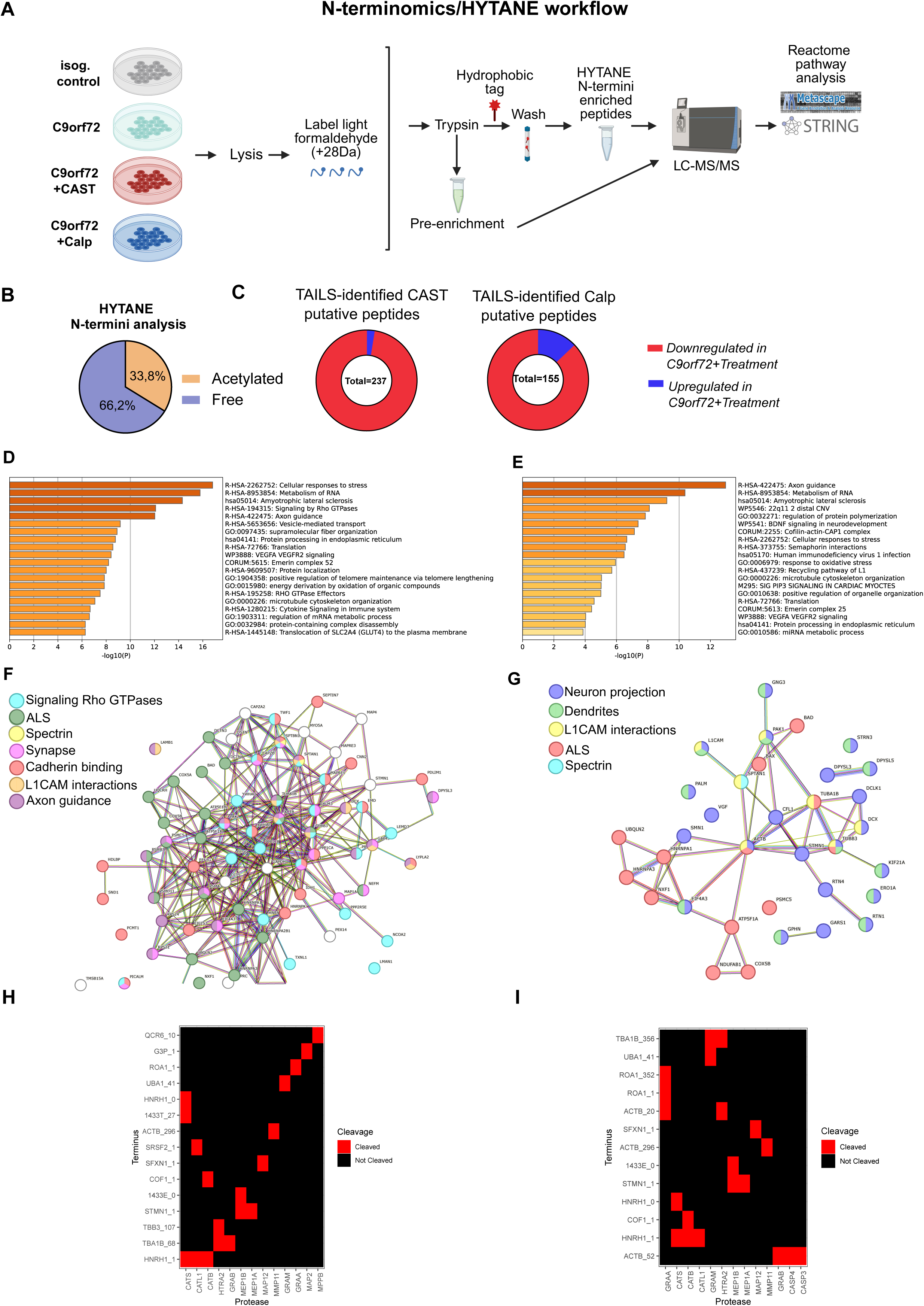
N-terminomics/HYTANE analysis of Isog. control and C9orf72 MN cultures identified synaptic substrates of CAST and Calptreatments. (**A**) Workflow schematic of N-terminomics experimental design. (**B**) Distribution of acetylated and free N-terminal peptides in the HYTANE enrichment. (**C**) Distribution of putative peptides showing downregulation (red) and upregulation (blue) following CAST and Calp treatments. (**D,E**) Pathway enrichment analysis. Bar plots showing significantly enriched pathways ranked by −log10(*P*). Terms are ordered by significance, highlighting the top biological processes associated with CAST (**D**) and Calp (**E**) treatment. (**F, G**) STRING-db (Szklarczyk et al. 2019) analysis of C9orf72 MNs treated with CAST (**F**) and Calp (**G**) showing enrichment for downregulated associated genes. For visualization purposes, only a selection of the associated genes is shown. (**H,I**) Protease cleavage heatmap for CAST (**H**) and Calp (**I**) treatment. Rows show protein termini and columns show proteases. Red indicates detected cleavage, while black indicates no cleavage.

Across all conditions, 33.8% of identified N-termini were acetylated and 66.2% were free N-termini, consistent with previous reports in cells ^44^ (**Fig 6B**). Comparative analyses between untreated and treated C9orf72 motor neurons enabled the identification of candidate CAST-and calpain-dependent cleavage sites. TopFIND-based annotation revealed numerous previously undescribed cleavage events (**Supp Tables 1 and 2**), substantially expanding the known repertoire of putative protease substrates in a disease-relevant neuronal context. N-terminal feature analysis further confirmed enrichment of canonical processing events, including methionine excision, signal peptide removal, and endoproteolytic cleavage.

Global quantitative analysis revealed a marked remodeling of the N-terminome upon treatment. The majority of proteins with detected N-termini exhibited reduced abundance in both CAST-treated (97.5%) and calpeptin-treated (87.1%) C9orf72 motor neurons, consistent with widespread proteolytic reprogramming (**Fig 6C**). Pathway enrichment analysis of treatment-dependent cleavage events (top 20 clusters; **Fig 6D,E**), using metascape, revealed strong overrepresentation of processes involved in neuronal development (including axon guidance and BDNF signaling), cytoskeletal dynamics, RNA metabolism and endoplasmic reticulum function. In addition, stress-response pathways including oxidative stress, autophagy, inflammatory and antiviral signaling, as well as insulin and VEGF pathways, were also prominently enriched, with ALS-related terms ranking consistently among the top categories.

Protein–protein interaction (PPI) network (STRINGdb enrichment analysis) (**Fig 6F, G, Supp Tables 3 and 4**) further showed that a substantial fraction of putative substrates localizes to synaptic compartments, including neuron projections, dendrites, and neuron–neuron synapses. This distribution aligns with the sites of functional rescue observed upon CAST and calpeptin treatment. CAST-modulated cleavage events were strongly enriched in synaptic signaling, cadherin binding, ALS-related KEGG pathways, and L1CAM, Rho GTPase, and axon guidance signaling (**Fig 6F**). Similarly, calpeptin-associated changes highlighted enrichment in neuronal projection and dendritic organization, L1CAM interactions, and ALS-related pathways (**Fig 6G**). Functional enrichment analysis using g: Profiler (**Table 1**) identified a highly overlapping set of genes involved in these same pathways, reinforcing the STRING results.

Together, these analyses reveal a shared functional landscape of proteins cleaved in C9orf72 MNs under both CAST and calpeptin treatments. Notably, the dataset also uncovered a broader spectrum of protease-associated signatures, with distinct cleavage patterns linked to multiple protease families, including GRAA, CATL1, MEP1B, MAP2, MMP1, and caspases such as CASP3 and CASP4 (**Fig 6H,I**). These findings suggest the activation of a complex and interconnected proteolytic network downstream of treatment.

Collectively, this HYTANE N-terminomics approach identifies ALS-relevant proteolytic events that are enriched in synaptic and axonal compartments, including previously reported calpain substrates. The results provide mew mechanistic insight into disease-associated proteolytic remodeling in C9orf72 ALS motor neurons.

### Calpastatin compensation attenuates the pathological accumulation of spectrin breakdown products in the axons of C9orf72 motor neurons

Interestingly, a subset of substrates was shared between the CAST and Calp treatment conditions, further supporting the robustness of the dataset (**Supp Table 5**). Among these, spectrin alpha (SPTAN1; Q13813) and spectrin beta (SPTBN1; Q01082) were particularly prominent. These cytoskeletal proteins are well known to be essential for axonal integrity, structural stability, and synaptic transmission and their proteolytic cleavage by caspases and calpains is well documented ^45–47^. The identification of spectrin alpha II (SPTAN1) as a substrate in both treatment conditions with its known sensitivity to proteolytic processing and reinforces the biological relevance and robustness of our N-terminomics results. To independently validate these findings, we examined spectrin processing in C9orf72 MNs. A disease-associated 150 kDa spectrin cleavage fragment was detected in C9orf72 cells but was absent in isogenic control cells (**Fig 7A**), indicating altered spectrin proteolysis in the context of C9orf72 ALS.

**Figure 7:**
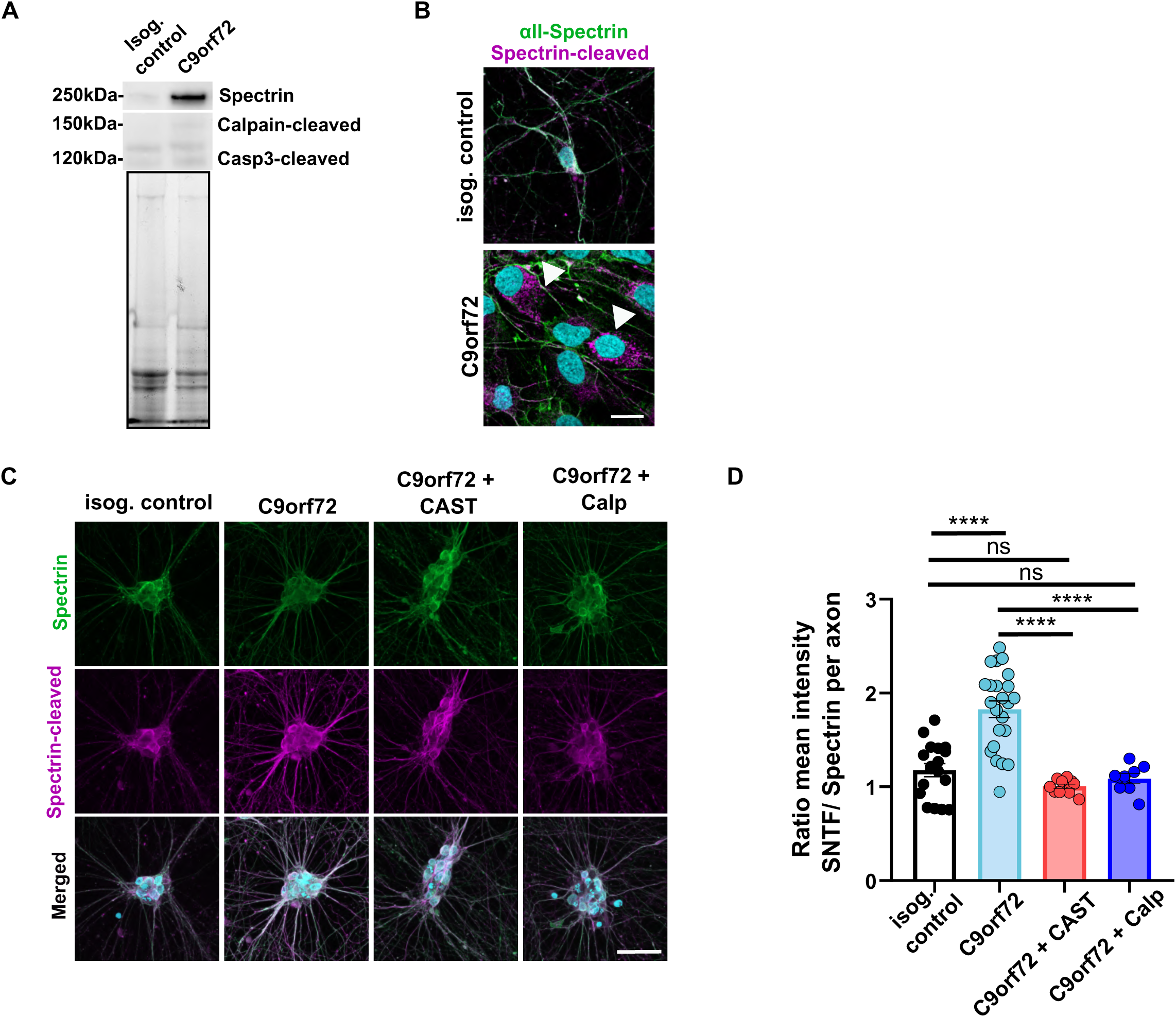
CAST and Calp reduce spectrin breakdown and SNTF accumulation in C9orf72 motor neurons. (**A**) Immunoblot of Spectrin levels indicate increased calpain cleaved 150-kDa spectrin breakdown product, and no change in 120-kDa caspase-3 cleaved product in C9orf72 MNs. Extractions were performed in MNs harvested after 3 weeks post-plating. (**B**) Representative immunostaining of Spectrin (green) and Spectrin-cleaved (SNTF) (purple) showing aggregates of spectrin breakdown product (arrowheads) in C9orf72 MN soma. Scale Bar: 10µm. (**C**) Representative images of Spectrin (green) and Spectrin-cleaved (purple) in isog. Control and C9orf72 MNs in comparison with C9orf72 treated MNs with either CAST or Calp, as indicated. Scale Bar: 50µm. (**D**) Quantification of the SNTF/spectrin ratio based on mean fluorescence intensity measured with Fiji, with analysis focused on axons (N=2, 1-way ANOVA). Data are represented as mean ± SEM. ns, not statistically significant; ****P<0.0001.

We next quantified the ratio of cleaved to full-length spectrin as a measure of proteolytic activity. Accumulation of spectrin N-terminal fragments (SNTF) was observed in the soma of C9orf72 motor neurons, suggesting impaired processing or clearance of cleavage products (**Fig 7B**). Importantly, targeted analysis of axons revealed a significant increase in the cleaved-to-full-length spectrin ratio under basal conditions, consistent with elevated axonal proteolytic activity (**Fig 7 C, D**). Remarkably, a 24-hour treatment with either CAST or Calp was sufficient to restore the cleaved-to-full-length spectrin ratio and its SNTF fragment to near-control levels (**Fig 7 C, D**). Together, these results corroborate our N-terminomics findings and strongly support the presence of a dysregulated proteolytic pathway in C9orf72 motor neurons that can be pharmacolically normalized. This normalization likely contributes to the preservation of neuronal integrity and function.

## DISCUSSION

Despite significant advances in ALS research and therapeutic development, no curative treatment is currently available ^20^. The existing approved therapies provide only marginal clinical benefit, underscoring the urgent need to identify novel therapeutic targets and compounds for treating ALS. In this study, we investigated the therapeutic potential of restoring calpastatin function, a key endogenous inhibitor of calpains, whose expression is markedly reduced in C9orf72 ALS models. Using two pharmacological agents, calpeptin and calpastatin peptide CAST, we show that both compounds significantly improve motor behavior and restore NMJ structure and function in the zebrafish C9orf72 ALS model. In C9orf72 MNs derived from patient IPSCs, pharmacological restoration of calpastatin function rescued neuronal death and normalized MN excitability. N-terminomics analysis revealed common substrate proteins modulated by both treatments including spectrin, highlighting a shared underlying mechanism and providing a rich dataset to identify additional synaptic substrates of proteases (calpain or perhaps others) regulated by calpastatin.

Our study also provides a more detailed characterization of the synaptic defects, both morphological and functional, in C9orf72 ALS models. Indeed, iPSC-derived MNs from a C9orf72 patient also exhibited presynaptic defects, such as enlarged synapsin-positive puncta, consistent with observations made in vivo in zebrafish model ^18^. In addition to altered synaptic transmission, we observed increased expression of the presynaptic protein synapsin in a sporadic ALS line, recapitulating previous findings in TDP-43 iPSC model ^35^ and FUS mouse models ^47^. Interestingly, although synapsin area was unchanged, its immunoreactivity was increased in FUS mice, potentially indicating a compensatory synaptic response.

Additionally, we identified a progression from early hyperexcitability to later hypoexcitability in these motor neurons, a pattern consistent with other ALS models, including mouse and iPSC-derived systems ^48–50^. Such changes in excitability may contribute to motor neuron vulnerability and degeneration by altering calcium homeostasis, increasing metabolic stress, and disrupting synaptic communication. Importantly, Calp and CAST restore synaptic integrity and neurotransmission at NMJs and synapses in both preclinical models.

The N-terminomics analysis provided a precise characterization of the proteomic changes induced by both CAST and Calp treatments, revealing significant enrichment of synaptic proteins. Among these, SPTAN1 (α-spectrin) emerged as a common substrate modulated by both treatments, confirming a shared mechanistic target. SPTAN1 has been implicated in a range of neurological disorders, including early-onset epileptic encephalopathy, developmental delay, and hereditary motor neuropathies. Cleavage of the spectrin cytoskeletal protein is a well-known calpain target critical for synapse stabilization ^51,52^, and selective inhibition with CAST or Calp mitigated the accumulation of cleaved spectrin in C9orf72 MN axons. N-terminomics further revealed modifications in both α-and β-spectrin, consistent with reduced proteolysis and restored overall abundance. These findings not only validate SPTAN1 as a key calpain substrate in C9orf72 ALS but also suggest that spectrin breakdown products (SBPs) could serve as potential biomarkers. This results align with previous studies reporting increased spectrin proteolysis in the hSOD1^G93A^ ALS mouse model ^14^, and in a traumatic brain injury model ^53^, where calpastatin overexpression reduced calpain-mediated spectrin cleavage and improved motor and cognitive behavioral outcomes. Moreover, spectrin breakdown products are widely recognized as reliable biomarkers of CNS injury, serving as powerful indicators of axon injury and degeneration ^54^. Our data therefore reinforce the concept that targeting the calpastatin-calpain pathway not only promotes motor neuron survival but also supports synaptic integrity, and provides candidate biomarkers to monitor therapeutic efficacy.

Given that synaptic dysfunction is a hallmark of ALS pathology ^55^, therapeutic strategies aimed at preserving neuromuscular synapses could prove highly beneficial. Under physiological conditions, calpains play important roles in neural development and synaptic transmission. However, during excitotoxicity, when intracellular calcium overload occurs, there is an excessive induction of calpain activation leading to deleterious effects ^56,57^. For example, calpain activation has been reported to trigger degradation of both pre and post-synaptic components ^58^. It has been shown that calpain can cleave synaptic proteins SNAP25 and SNAP23 impairing synaptic vesicle fusion and exocytosis ^59–64^. Moreover, upon intracellular Ca^2+^ accumulation, calpain activity results in cleavage of ALS proteins TDP-43, C9orf72, matrin3, VCP, profilin-1 and FUS ^65^. We demonstrated a neuroprotective effect of the inhibition of the calpain activity with its inhibitor calpeptin or by restoring calpastatin function with the active peptide CAST on motor behaviour and NMJ function in zebrafish C9-miR ALS model.

We demonstrated that restoring calpastatin function in C9-miR zebrafish with Calp or CAST preserved quantal synaptic transmission and synaptic vesicle cycle at NMJ. Dysfunctional NMJ transmission occurs at early stages in C9orf72 ALS ^48,66^. Our findings suggest that enhancing calpastatin function may prevent NMJ dysfunction in ALS and represents a promising therapeutic avenue for further investigation. Furthermore, reduced expression of the synaptic protein SV2A has been reported in cortical and MNs derived from C9orf72 patient induced pluripotent stem cell lines ^67^ and in the zebrafish C9-miR ALS model ^18^. Expression of human calpastatin in a mouse model of SCS increases SV2A levels and significantly rescues impaired neuromuscular transmission at NMJ ^17,68^. Alterations in the calpastatin/calpain system have been shown to activate Cdk5 ^69^, negative regulator of synaptic vesicle recycling and can repress presynaptic neurotransmission ^70–72^. Overexpression of a Cdk5 inhibitory peptide in MNs of SOD1^G37R^ ALS mice improves motor deficits, extends survival and delays neuroinflammation and pathology in brain and spinal cord of these mice. Thus, a thorough characterization of the calpastatin/calpain/Cdk5 cascade may provide new insights into synaptic dysfunction in ALS and uncover a broad range of potential therapeutic target.

Calpain inhibition shows considerable promise as a neuroprotective strategy in ALS. Previous studies strongly support this role ^73^. Selective calpain inhibitors have been shown to be neuroprotective in the hSOD1^G93A^ ALS mouse model ^14^, extending survival by 63 days and delaying the motor axon degeneration. Additionally, calpastatin overexpression reduces toxicity of SOD1^G93A^ in dissociated spinal cord cultures, thereby prolonging culture viability ^74^. Furthermore, Yamashita et al. demonstrated that phosphorylated TDP-43 is more resistant to calpain-dependent cleavage *in vitro* compared to its non-phosphorylated counterpart, underscoring the pivotal role of calpain in TDP-43 pathology and aggregation. Here, we provide the first evidence that targeting calpastatin/calpain signaling mitigates disease phenotypes in both sporadic and C9orf72 ALS models, using zebrafish and human iPSC-derived MNs. Pharmacologically mimicking calpastatin function with Calp or CAST improves motor behavior, NMJ function, and synaptic integrity, supporting their potential as broad ALS therapeutics. Calpastatin neuroprotective effects have been reported in several neurodegenerative diseases. For instance, in Huntington’s disease, preventing calpastatin degradation and stabilizing its function with the drug CHIR99201 reduced neuropathology and ameliorated behavioral deficits mice models ^75,76^. The same treatment improved mitochondrial function and promoted neuronal survival in Huntington’s disease patient-derived neurons ^75^. Multiple studies further suggest that excessive calpain activity contributes to spinal cord degeneration and pathogenesis in Parkinson’s disease (PD) ^77–79^. Accordingly, calpain inhibition has been shown to improve behavioral outcomes in MPTP-induced PD models ^78^.

## CONCLUSION

In summary, our study demonstrates that restoring calpastatin function preserves NMJ integrity, maintains synaptic architecture, and improves motor neuron survival in *in vivo* and *in vitro* ALS models. N-terminomics approach not only identified key substrates such as SPTAN1 but also provides a rich resource for uncovering additional mechanistic pathways and potential biomarkers. These findings highlight the therapeutic potential of targeting calpastatin-dependent pathways in both C9orf72 familial and sporadic ALS.

## METHODS

### Zebrafish Husbandry

Wild-type and C9-miR adult zebrafish (*Danio rerio*) were maintained at 28°C on a 12/12 h light/dark cycle in accordance with The Zebrafish book ^80^. The C9-miR (c9orf72 knockdown) zebrafish line was generated using a miRNA-based gene-silencing approach^18^.Embryos were raised at 28.5 °C and collected and staged as previously described ^81^.

### Gene expression study

RNA was isolated from approximatively 30 larvae (6 dpf) using TriReagent® (Sigma Aldrich) according to manufacturer’s protocol. 1µg of RNA was used for cDNA synthesis using the SuperScript®Vilo™ kit (Invitrogen). Real-time qPCR (RT-qPCR) was run with SYBR Green Master Mix (Bioline) using the LightCycler® 96 (Roche). *ef1a* was used as the reference gene for normalization and following primers were used for *cast*: *cast* Forward primer: 5’-ACAGACAAGTGCTCAAAAGGTG-3’; Reverse primer: 5’ - GATCGTCAGCATCTGCACTG-3’.

### Zebrafish drug treatment

Zebrafish embryos at 48 hours post fertilization were placed in petri dishes and treated for 4 days in embryo medium (E3). Calpeptin (Sigma, #C8999) was used at 10μM, stock was reconstituted in DMSO. Calpastatin peptide (Sigma-Aldrich, #SCP0063) was used at 1µM, stock was reconstituted in distilled water. Both drugs and water were changed every 48 hours.

### Behavioural assay

Larvae (6 dpf) were transferred into single wells of a 96-well plate containing 200µl of E3 media and habituated in the the Daniovision® recording chamber (Noldus) for 1 hour before starting the experiment. Larval locomotor activity was examined using the Daniovision® apparatus. Analysis was performed using the Ethovision XT12 software (Noldus) to quantify the total swimming distance in given hours and the locomotor activity per second.

### NMJ morphology in larval zebrafish

Immunohistochemical analyses were performed on 6 dpf zebrafish to visualize NMJ pre-and postsynaptic structures. Briefly, animals were fixed in 4% paraformaldehyde overnight at 4°C. After fixation, the larvae were rinsed several times for 1 hour with PBS-Tween and then incubated in PBS containing 1 mg/ml collagenase for 180 minutes to remove the skin. The collagenase was washed off with PBS-Tween for 1 hour, and larvae were then incubated in blocking solution (2% Normal Goat serum (NGS), 1% Bovine Serum Albumin (BSA), 1% DMSO, 1% Triton-X in PBS (PBST)) containing 10 mg/ml Tetramethyl rhodamine-conjugated α-bungarotoxin (Thermo Fisher Scientific, Cat#T1175) for 30 minutes. The larvae were rinsed with PBST and then incubated in freshly prepared blocking solution containing primary antibody anti-Synaptic vesicle glycoprotein 2A (SV2A) (Developmental Studies Hybridoma Bank, Cat#AB_2315387) overnight at 4°C. The next day, larvae were incubated in blocking solution containing a secondary antibody (Alexa Fluor™ 488, 1:1000, Invitrogen™, Cat#A-21042) overnight at 4°C. The larvae were then washed with PBST and mounted on a glass slide in 80% glycerol. Z-stack imaging was performed using confocal laser scanning microscope model LSM780 (Carl Zeiss) and images were analyzed with ZEN software (Carl Zeiss). Co-localization analysis of pre-and post-synaptic structures was performed using the JACOP program from ImageJ (NIH).

### FM1-43 staining

Zebrafish larvae (6dpf) were first anesthetized in Evan’s solution (134 mM NaCl, 2.9 mM KCl, 2.1 mM CaCl_2_, 1.2 MgCl_2_, 10 mM HEPES, 10 mM glucose) containing 0.02% tricaine (Sigma Aldrich). The larvae were then pinned to a Sylgard coated dish both at the head and extreme tail end using electrolytically sharpened tungsten needles. The skin was then carefully peeled away to expose the muscles and to permit access to FM1-43 (Molecular Probes, #T3163). The fish were treated with Evan’s solution containing 10 μM of FM1-43 to allow preloading penetration of the dye molecules. After 10 min, the fish were transferred to a high potassium Hank Buffer Salt Solution (HBSS) (97 mM NaCl, 45 mM KCl, 1 mM MgSO_4_, 5 mM HEPES, 5 mM CaCl_2_) containing 10 μM of FM1-43 for 5 minutes. The fish were then transferred in Evan’s solution with 10 μM of FM1-43 finished for an additional 3 minutes, after which loading was complete. The fish were then washed with a low calcium Evan’s solution (0.5 mM CaCl_2_) three times for 5 minutes to minimize spontaneous release of loaded synaptic vesicles. The fish were imaged for FM1-43 staining using a 40X Examiner A1 microscope (Zeiss). Blind measurements of FM1-43 staining at NMJs in wild-type control and C9-miR fish were performed using ImageJ (NIH).

### Electrophysiology recordings

Whole-cell patch-clamp recordings were made from muscle cells of 6 dpf larvae. The preparation was bathed in an extracellular solution (134 mM NaCl, 2.9 mM KCl, 1.2 mM MgCl, 10 mM HEPES, 10 mM glucose, pH 7.8) containing 1 μM tetrodotoxin (TTX; Tocris, UK) to block action potentials during recordings of miniature endplate currents (mEPCs). Patch-clamp electrodes (2-4 MΩ) were filled with an intracellular solution (130 mM CsCl, 8 mM NaCl, 2 mM CaCl2, 10 mM HEPES, 10 mM EGTA, 4 Mg-ATP, 0.4 Li-GTP, pH 7.4). Miniature endplate currents from white muscle fibres were recorded in whole-cell configuration with an Axopatch 200B amplifier (Molecular Devices) at a holding potential of-60 mV, low-pass filtered at 5 kHz and digitised at 50 kHz. Series resistance was compensated for by at least 85% using the amplifier’s compensation circuitry. Synaptic currents were recorded using pCLAMP10 software (Molecular Devices).

### Analysis of miniature endplate currents

mEPCs were analysed using AxoGraph X software. The mEPC recordings were examined by the software and synaptic events were detected using a template function. Overlapping or misshapen events were removed, and the remaining events were averaged, and the properties (amplitudes, decay time constants, frequencies) of the averaged trace were measured. Events with slow rise times and low amplitudes were excluded from the analysis, so that only events with fast rise times were included in our analysis, as these events originated from the patch-clamped cells rather than from nearby electrically coupled muscles. Individual decay time constants were fitted over the initial (fast) decay portion and over the distal (slow) portion of the decay. For each *n*, currents were recorded from a single white muscle fibre from a single larva.

### CRISPR editing

We generated an isogenic control iPSC line by CRISPR/Cas9-mediated genome editing. The patient-derived iPSCs were nucleofected with a combination of Cas9 protein and two sgRNAs targeting the *C9orf72* gene. sgRNA1 and sgRNA2 were designed to create two double-strand breaks (DSBs) flanking the G4C2 hexanucleotide repeat, resulting in deletion of the pathogenic expansion. sgRNA1 was located 141 bp upstream and sgRNA2 108 bp downstream of the repeat. Following single-clone selection, edited clones were screened using ddPCR and Sanger sequencing, confirming biallelic deletion of the G4C2 repeat. The resulting clone carried a 261 bp deletion corresponding to the flanking region of the expansion.

To generate the isogenic control line, the edited clone was corrected using a donor template. This template, synthesized by Bio Basic and cloned into the pUC57 plasmid, was nucleofected along with sgRNA3 and Cas9 protein. After clonal selection, edited clones were analyzed by Sanger sequencing and Repeat-Primed PCR (AmplideX® PCR/CE C9orf72 Kit), confirming the integration of the 261 bp fragment into both alleles. The resulting isogenic control line carried three G4C2 repeats (**Fig Supp 1C**).

### CRISPR/Cas9

CAS9 protein (1 μl; stock 61 μM), sgRNAs (3 μl; stock 100 μM) and ssODNs (1 μl; stock 100 μM) in 20 μl of nucleofection buffer P3 were nucleofected (program CA137, 4D-Nucleofector Device, Lonza) into 500,000 detached iPSCs ^23^. After single clone selection, gene-edited clones were identified by ddPCR (QX200™ Droplet Reader, Bio-Rad) Sanger sequencing and Repeat-Primed PCR (AmplideX® PCR/CE C9orf72 Kit).

### iPSC lines and reprogramming

PBMCs were obtained through the C-BIG Repository at the Montreal Neurological Institute (The Neuro). For lines used in this study included the control cell line AIW002-02 (C-BIG ID#: IPSC0063-male, 37 years old), the C9orf72 ALS patient cell line 3414 ((C-BIG ID#: IPSC0049-male, 66 years old) paired with its isogenic correction and a sporadic ALS patient cell line TD17 ((C-BIG ID#: IPSC0092-Male, 49 years old). These lines were all reprogrammed from peripheral blood mononuclear cells (PBMCs) and underwent the QC profiling as described in earlier work ^82^. The C9orf72 iPSC line was generated in-house by episomal reprogramming ^83^. The expansion mutation was validated to be higher than 145 G4C2 repeats through Sanger sequencing and Repeat-Primed PCR (AmplideX® PCR/CE C9orf72 Kit). Briefly, PBMCs were cultured for 6 days, 2∼3 x10^6^ cells were nucleofected with episomal plasmids (pEV-OCT4-2A-SOX2, pEV-MYC, pEV-KLF4, and pEV-BC-XL, a generous gift from Dr. XB Zhang, Loma Linda University). The transfected PBMCs were plated on mitomycin C-treated mouse embryonic fibroblasts cultured in KnockOut DMEM/F12 supplement with 10% Knockout serum supplement, 50 ng/ml fibroblast growth factor 2, 1x Insulin-Transferrin-Selenium and 50 mg/ml 2-phospho-L-ascorbic acid. The cultures were refreshed with 2 ml of the above medium every 2 days until day 8. When colonies displayed iPSC morphology (between day 6-8 post-transfection), cells were fed with mTeSR1 medium (Stemcell Technologies) supplemented with 0.25 mM sodium butyrate every 2 days until day 14. Colonies were picked manually on day 14-16 and cultured on Matrigel-coated dishes every 5–7 days until after 5 passages when they were cryopreserved for further testing and profiling. Each experiment was performed using technical replicates generated by independent differentiations.

### Culture conditions for iPSCs

iPSCs were cultured and expanded on plates coated with Matrigel (Corning Millipore; Cat#354277) in E8 (ThermoFisher Scientific, Cat#A1517001) media. Cells were maintained at 37°C with 5% CO2 with daily media changes and split when cells reached 70-80% confluency (within 5-7 days of seeding). Any iPSC colonies with irregular borders, spontaneous differentiation or transparent centers were manually removed prior to splitting. Cells were passaged by incubation in Gentle Cell Dissociation Reagent (Stemcell Technologies; Cat#07174) for 6 minutes at room temperature to obtain small aggregates of colonies. iPSC lines were first differentiated into human MNs progenitor cells (MNPCs) on dishes coated with 10 μg/ml PLO and 5 μg/ml laminin (Life Technologies; Cat#23017-015) and finally MNs according to a previously published protocol ^22,35^. The following cell densities were used: 5×10^4^ cells/well in 24 well plates for immunocytochemistry; 2×10^5^/60 mm dish for gDNA extraction.

### Karyotyping and genomic abnormalities analysis

Genomic DNA was extracted with the Genomic DNA Mini Kit. Genomic integrity was detected with the hPSC Genetic Analysis Kit (Stemcell Technologies; Cat#07550) according to the manufacturer’s instructions. Briefly, 5 ng of genomic DNA was mixed with a ROX reference dye and double-quenched probes tagged with 5-FAM. The probes represented eight common karyotypic abnormalities that have been reported to arise in hiPSCs: chr 1q, chr 8q, chr 10p, chr 12p, chr 17q, chr 18q, chr 20q or chr Xp (**Fig Supp. 2B-C)**. Sample-probe mixes were analyzed on a QuantStudio 5 Real-Time PCR System (ThermoFisher Scientific). Copy numbers were analyzed using the ΔΔCt method. The results were normalized to the copy number of a control region in chr 4p ^84^. For G-band karyotyping, iPSCs were cultured for 72 hours until they attained 50-60% confluency, then were shipped live to the Centre for Applied Genomic, The Hospital for Sick Children (Toronto, ON).

### Viability assay

MNPCs were seeded at a density of 15,000 cells per well in opaque white optical 96-well plates coated with PLO/laminin. The cells were cultured in the final differentiation medium for one week before being treated with 1mM cytosine arabinoside (AraC, Sigma-Aldrich; Cat#C6645) for 6 hours to eliminate any residual proliferating cells. After 10 days, media was fully replaced with fresh final differentiation medium or with fresh final differentiation medium containing half the concentration of N2 (Life Technologies; Cat#17502–048) and B27 (Life Technologies; Cat#17504–044) supplements, without neurotrophic factors (i.e., BDNF (Peprotech; Cat#450–02), CNTF (Peprotech; Cat#450–13), and IGF-1 (Peprotech; Cat#100–11) as previously described ^35^. At four-weeks post-plating (i.e., 28 days of differentiation), MNs were exposed to 5µM active CAST peptide or 10µM calpeptin for 24 hours. The viability of MNs was then assessed using an ATP-based chemiluminescent assay (Cell Titer-Glo, Promega, #G7570) following the manufacturer’s instructions and the luminescence of the neuronal cultures was acquired with a GloMax Microplate Reader (Promega). The percentage of viability of MNs was determined by normalizing the raw luminescence values to those of the 1-week reading (where viability was assumed to be 100%) of each cell line to account for differences in plating.

### Quantitative PCR

Total RNA was isolated from differentiated MNs at 1-, 3-, 4-and 6-weeks post-plating uusing TriReagent® (Sigma) according to manufacturer’s protocol. 1µg of purified RNA was used for cDNA synthesis using the SuperScript®Vilo™ kit (Invitrogen; Cat#11754050). RT-qPCR reactions were run in triplicates with SYBR Green Master Mix (Biorad) using the LightCycler® 96 (Roche). GAPDH was used as the reference gene for normalization and following primers were used for human CAST: Forward primer: 5’-CAAAAAGCCTACCCAAGCAG 3’; Reverse primer: 5’ GCACAGCTGGGGTTAATGAT 3’.

### Immunofluorescence staining

For immunofluorescence staining, the cells were plated on 12-mm circular coverslips, fixed in 4% formaldehyde for 20 min at room temperature and washed three times with PBS.

Cells were then permeabilized with 0.2% Triton X-100 in PBS for 10 min and blocked for 1 hour at room temperature in blocking solution (5% normal donkey serum (NDS, Millipore; Cat#S30-100), 1% bovine serum albumin (BSA, Multicell; Cat#800–095-CG) and 0.05% Triton X-100 in PBS). Cells were incubated with primary antibodies in blocking buffer overnight at 4 °C. After three washes in PBS, cells were incubated with appropriate secondary antibodies for 2 hours at room temperature, followed by staining with Hoechst DNA dye (Life Technologies; Cat#H3570) for 10 min. Fluorescence images were captured using a confocal laser scanning microscope, model LSM700 (Carl Zeiss) and automated Evos FL-Auto2 imaging system (ThermoFisher Scientific). The following primary antibodies were used: anti-NF-H (EnCor Biotech; Cat#CPCA-NF-H), anti-calpastatin (Thermo Fisher Scientific; Cat#1F7E3D10), anti-Synapsin I (Millipore; Cat#574777), anti-PSD95 (Millipore; Cat#MABN68), anti-α-II-Spectrin (BioLegend; Cat#803201), anti-SNTF (Millipore; Cat#ABN2264).

### Western blot

Proteins were collected from differentiated MNs at 1-, 3-, 4-and 6-weeks post-plating. For Western blotting, 20 μg of protein per sample in Laemmli buffer was resolved on 7.5% or 10% SDS-PAGE gels and transferred to PVDF or nitrocellulose membranes using the Trans-Blot Turbo Transfer System (Bio-Rad). The membranes were blocked for 1 hour with 5% non-fat milk solution in 1× phosphate buffered saline (PBS). Primary antibodies were then applied in blocking solution and incubated overnight at 4°C. The following primary antibodies were used: (anti-calpastatin (Thermo Fisher Scientific; Cat#1F7E3D10), anti-calpain-2 (Cell Signaling; Cat#70655), anti-Synapsin I (Calbiochem; Cat#574777), anti-α-II-Spectrin (BioLegend; Cat#803201), anti-SNTF (Millipore; Cat#ABN2264). After three washes with TBS-Tween (0.1%), membranes were incubated for 2 hours at room temperature with horseradish peroxidase (HRP)-conjugated secondary antibodies (1:8000 dilution) in blocking solution. The following HRP-conjugated secondary antibodies were used: Anti-Mouse (Jackson Immunoresearch, Cat#115-035-003), Anti-Rabbit (Jackson Immunoresearch, Cat#111-035-144). Bands were visualized Clarity Western ECL Substrate or Clarity Max Western ECL Substrate (Bio-Rad; Cat#170–5061, Cat#1705062) and imaged using the ChemiDoc MP Imaging System (Bio-Rad).

### Microelectrode array (MEA) recordings

Cytoview 24-well MEA plates (#M384-Tmea-24w, Axion Biosystems) were coated with a 10 μg/mL solution of poly-L-ornithine (PLO, #P3655, Sigma-Aldrich) diluted PBS. After 24 hours of incubation at 37 °C, the PLO solution was removed by performing three washes with 1X PBS. Immediately after, the MEA plates were coated with a 5 μg/mL solution of laminin (Sigma-Aldrich; Cat#L2020) diluted in DMEM/F-12 (Thermo Fisher Scientific; Cat#10565018) and incubated for another 24 hours at 37 °C.

50,000 MNPCs were plated in final MN differentiation medium onto the electrode area of the MEA 24-well plates and MEA recordings were conducted at 1-, 2-, 3-, 4-, 5-, and 6-weeks post-plating. Before each recording, the MN final differentiation medium was replaced with freshly prepared sterile 1X artificial cerebrospinal fluid (aCSF) (osmolarity 305mOsm/L and pH 7.4), as previously described ^85^. MEA plates containing MNs cultures in 1X aCSF were incubated for 1 hour before being transferred to the Axion Maestro Edge (Axion Biosystems) where they were then allowed to equilibrate at 37 °C and 5% CO₂ for approximately 5 minutes prior to recording. Data were collected for 5 minutes using the Axis Navigator software (Axion Biosystems, version 1.5.1.12, Atlanta, GA, USA), with a band-pass filter set from 3 kHz (low-pass) to 200 Hz (high-pass).

Following the recording, the MEA plates were removed from the instrument, and the 1X aCSF was replaced with MN final differentiation medium to maintain the cells in culture for future recording sessions.

### Proteomics sample preparation

Isogenic control and C9orf72 ALS MNs were lysed using 1 mL lysis buffer (1% SDS, 10 mM EDTA, 100 mM ammonium bicarbonate and 1x protease inhibitor tablets (Roche complete) in high-performance liquid chromatography water (HPLC water), pH 8.0) and mechanically scraped. Cell contents were collected in a 2 mL Eppendorf tube and sonicated to shear DNA. Lysates were then centrifuged at 14,000g for 10 min in a microcentrifuge pre-cooled to 4°C to pellet DNA. The supernatant was collected, and the protein concentration was measured using a BCA assay according to manufactures directions (Thermo Fisher Scientific; Cat#23225).

### N-terminomics/HYTANE and shotgun proteomics

Isogenic control/ C9orf72-drug/ C9orf72+Calp/ C9orf72+CAST cells were treated with PMA overnight followed by a 1 h treatment with the calcium ionophore (A23187; Sigma; Cat#C7522). Samples were lysed (1% SDS, 10 mM EDTA, 100 mM ammonium bicarbonate and 1x protease inhibitor tablets (Roche complete) in high-performance liquid chromatography water (HPLC water), pH 8.0) and subjected to an N-terminomics/HYTANE workflow (**Fig 6A**) ^38–40,42,86–91^. 400 ug of samples was diluted in 3 M final guanidine hydrochloride. Samples were reduced with 25 mM DTT (Gold Biotechnology, St. Louis, MO) at 37°C for 1 h and alkylated with 15 mM IAA (GE Healthcare, Mississauga, ON) in the dark at room temperature for 30 min followed by quenching with 15 mM DTT. The pH was adjusted to 6.5 before the samples were labeled with a final concentration of 40 mM light formaldehyde (VWR; Cat#10790-709) in the presence of 40 mM sodium cyanoborohydride (Sigma; Cat#156159-10g) overnight at 37°C. Next, samples and were precipitated using acetone/methanol (8:1) overnight at - 20°C. Proteins were pelleted by centrifuging at 9 000 xg for 15 minutes at 4 °C. The proteins were washed in 100% methanol 3 times using the same centrifuge settings. The resulting pellet was resuspended by shaking in 200 ul of 1M NaOH, for 15 minutes. The pH was adjusted by adding 200 ul of 200 mM HEPES buffer before adjusting the pH to 8. Proteins were digested with 25 ug of trypsin (Thermo Fisher Scientific; Cat#90058) overnight at 37°C. For pre-enrichment HYTANE (pre-HYTANE)/shotgun proteomics, 10% of the trypsin digested samples were collected and the pH was adjusted to 3 with 100% Trifluoroacetic acid (TFA) and stored at-20°C. The remaining samples were adjusted to a pH of 6.5. 100% ethanol was added to a final concentration of 40% v/v. Samples were incubated for 1 hour at 37°C with 5% v/v of undecanal, and 40 mM sodium cyanoborohydride to complete the undecanal tagging of neo N-termini ^92^. Undecanal tagged peptides were removed by Solid Phase Extraction by washing on c18 columns (waters WAT036810) preconditioned with 95% methanol. Samples were lyophilized in a SpeedVac and resuspended with 1% TFA by shaking. Both pre-HYTANE and HYTANE samples were then desalted using Sep-Pak C18 columns and lyophilized before submitting for LC–MS/MS analysis to the Southern Alberta Mass Spectrometry core facility, University of Calgary, Canada.

### High performance liquid chromatography (HPLC) and mass spectrometry (MS)

Using a process previously described by Gordon et al. 2019 ^93^ tryptic peptides were analyzed on an Orbitrap Fusion Lumos Tribrid mass spectrometer (Thermo Scientific) operated with Xcalibur (version 4.4.16.14) and coupled to a Thermo Scientific Easy-nLC (nanoflow Liquid Chromatography) 1200 system. A total mass of 2 μg tryptic peptides were loaded onto a C18 trap (75 um x 2 cm; Acclaim PepMap 100, P/N 164946; ThermoScientific) at a flow rate of 2ul/min of solvent A (0.1% formic acid in LC-MS grade water). Peptides were eluted using a 120 min gradient from 5 to 40% (5% to 28% in 105 min followed by an increase to 40% B in 15 min) of solvent B (0.1% formic acid in 80% LC-MS grade acetonitrile) at a flow rate of 0.3 μL/min and separated on a C18 analytical column (75 um x 50 cm; PepMap RSLC C18; P/N ES803; ThermoScientific). Peptides were then electrosprayed using 2.1 kV voltage into the ion transfer tube (300°C) of the Orbitrap Lumos operating in positive mode. The Orbitrap first performed a full MS scan at a resolution of 120000 FWHM to detect the precursor ion having a m/z between 375 and 1575 and a +2 to +7 charge. The Orbitrap AGC (Auto Gain Control) and the maximum injection time were set at 4e5 and 50 ms, respectively. The Orbitrap was operated using the top speed mode with a 3 sec cycle time for precursor selection. The most intense precursor ions presenting a peptidic isotopic profile and having an intensity threshold of at least 5000 were isolated using the quadrupole and fragmented with HCD (30% collision energy) in the ion routing multipole. The fragment ions (MS2) were analyzed in the ion trap at a rapid scan rate. The AGC and the maximum injection time were set at 1e4 and 35 ms, respectively, for the ion trap. Dynamic exclusion was enabled for 45 sec to avoid of the acquisition of same precursor ion having a similar m/z (plus or minus 10 ppm). The mass spectrometry proteomics data have been deposited to the ProteomeXchange Consortium via the PRIDE ^94^ partner repository with the dataset identifier PXD067134.

### Proteomic data and bioinformatic analysis

Spectral data were matched to peptide sequences in the human UniProt protein database (version UP000005640 containing 83,385 entries and 20,644 genes) using the MaxQuant software package v2.5.2.0, peptide-spectrum match false discovery rate (FDR) of <0.01 for the preHYTANE proteomics data and <0.05 for the N-terminomics/Hytane data. Search parameters included a mass tolerance of 20 ppm for the parent ion, 0.05 Da for the fragment ion, carbamidomethylation of cysteine residues (+57.021464), variable N-terminal modification by acetylation (+42.010565 Da), and variable methionine oxidation (+15.994915 Da), Label setting included a light formaldehyde modification (+28.031300), and LFQ was turned on. For the N-terminomics/HYTANE data, the cleavage site specificity was set to semi specific free N-term ArgC (search for free N-terminus) for the HYTANE data and was set to semi specific ArgC for the preHYTANE data, with up to two missed cleavages allowed. Significant outlier cut-off values were determined after log(2) transformation by boxplot-and-whiskers analysis using the BoxPlotR tool ^95^.

Database searches were limited to a maximal Size of 6600 Da per peptide. Peptide sequences matching reverse entries were removed.

### Reactome pathway analysis

To identify interconnectivity among proteins, the STRING-db (Search Tool for the Retrieval of Interacting Genes) database was used to identify interconnectivity among proteins. The protein–protein interactions (PPI) are encoded into networks in the STRING.v11 ^96^ database (https://string-db.org). Our data were analyzed using Homo sapiens as our model organism at a false discovery rate of 1%.

## Statistics

For zebrafish experiments, N refers to a single egg clutch obtained from one mating, and n denotes the sample size. For iPSC-derived motor neuron experiments, N corresponds to an independent differentiation, while n represents the sample size (independent wells). Significance was determined using either Student’s t-test or One-way ANOVA followed by multiple comparisons test. A Tukey post-hoc multiple comparisons test was used for normally distributed and equal variance data. Kruskal-Walllis ANOVA and Dunn’s method of comparison were used for non-normal distributions. All graphs were plotted using the Graphpad PRISM software.

## Declarations Study approval

All the animal experiments were performed in compliance with the guidelines of the Canadian Council for Animal Care and received approval from the INRS ethics committee. The use of human cells in this study was approved by McGill University Research Ethics Board (IRB Study Number A03-M19-22A and DURCAN_iPSC/2019–5374).

## Availability of data and materials

All data associated with this study are present in the paper or the Supplementary Materials. Proteomic data are available via ProteomeXchange with identifier PXD067134.

## Author contributions

Conceptualization: LL, MC, TMD, and SP. Methodology: LL, ZB, CZ, MC, DY, MN, CQC, MN, ZY, NA and SP. Investigation: LL, ZB, CZ, MC, GH and SP. Visualization: LL and ZB. Funding acquisition: TMD, AD and SP. Project administration: TMD, AD and SP. Supervision: TMD, AD and SP. Writing—original draft: LL, ZB, and SP. Writing—review and editing: LL, ZB, CZ, DY, MC, TMD, AD and SP.

## Acknowledgments

The authors would like to thank Leonid Saponiv for animal care and husbandry.

LL is supported by a FRQS Postdoctoral Fellowship https://doi.org/10.69777/374842. SP is supported by the Anna Sforza Djoukhadjian Research Chair in ALS from Fondation Armand-Frappier, Natural Science and Engineering Research Council (NSERC), Canadian Institutes of Health Research (grant number 204072), and a FRQS Junior 2 research scholar. TMD is supported by the Canada First Research Excellence Fund, awarded through the Healthy Brains, Healthy Lives initiative at McGill University; the CQDM FACs program; and the Canadian Foundation for Infrastructure (CFI) funded JELF program. AD is supported by supported by the STARS award by the Arthritis Society of Canada, the Canadian Institutes of Health Research (CIHR) (grant number 449589) and the Natural Science and Engineering Research Council of Canada (NSERC) (grant number DGECR-2019-00112).

## Consent for publication

Not applicable.

## Ethics approval and consent to participate

Adult and larval zebrafish (*Danio rerio*) were maintained and experimental procedures were performed in compliance with the Canadian Council on Animal Care guidelines and approved by the INRS-LNBE ethics committee. All procedures with human material were approved by the Ethical Committees in Human Research of McGill University (DURCAN_iPSC/2019–5374 and IRB Study Number A03-M19-22A).

---

**Table 1.** List of CAST and Calp treatment-dependent cleavage events identified in C9orf72 MNs and their associated enriched-pathways.

