## Supplemental Material for "Targeting calpastatin pharmacologically restores synaptic proteolysis and preserves motor neurons survival and function in C9orf72 ALS"

**Short Title:** Targeting calpastatin rescues C9 ALS defects

**Supplemental Figure Legends**

**Fig Supp 1**. **Generation of the *C9Orf72* isogenic control iPSC line. (A)** Schematic representation of the strategy for the generation of the C9Orf72 isogenic control line by CRISPR/Cas9-mediated genome editing. **(B)** AmplideX-C9 PCR/CE profiles of the C9Orf72 patient's line, carrying 8 repeats ($G_4C_2$) on one allele and >145 on the other one (top) and the C9Orf72 isogenic control line carrying 3 repeats ($G_4C_2$) on both alleles (bottom). **(C)** C9Orf72 isogenic control iPSC line genotyping using Sanger sequencing.

**Fig Supp 2**. **Characterization of *C9Orf72* and *C9Orf72* isogenic control iPSC lines. (A)** Representative images of *C9Orf72* and *C9Orf72* isogenic control iPSCs subjected to immunocytochemistry for pluripotency associated markers SSEA-4, OCT-3/4, Nanog and TRA1-60. Scale bar, 200 μm. **(B and C)** Genomic stability analyses. *C9Orf72* and *C9Orf72* isogenic control iPSCs lines display normal G-band karyotypes **(B)** and have normal chromosome copy numbers, as assessed by qPCR. Data shown as mean ± SEM of technical triplicates. The control used here was provided by the manufacturer of the genome stability testing kit **(C)**.

**Fig Supp 3**. **Characterization of iPSC-derived MNPCs and MNs. (A)** Representative images of *C9Orf72* isogenic and *C9Orf72* MNPCs differentiated from iPSCs subjected to immunocytochemistry for the common MNPC marker Nestin and the proliferative marker Ki67. Nuclei were counterstained with Hoechst 33342. Scale bar, 100 μm. **(B)** Normalized expression levels of NANOG, POU5F1, OLIG2 and MNX1 in iPSCs, MNPCs, and iPSC-derived MNs differentiated for 14 and 28 days from *C9Orf72* isogenic (top panel) and *C9Orf72* (bottom panel) lines. Data normalized to Actβ-GAPDH expression. Bar graphs show the mean (SD).

**Fig Supp 4. Cellular distribution of calpastatin is altered in *C9Orf72* MNs. (A)** Representative images of MNs differentiated for 4 weeks stained for Calpastatin (green), Calpain2 (grey), and Neurofilament Heavy Chain NF-H (red). Nuclei were counterstained with Hoechst (blue). Scale bar: 50µm. (**B**) Bar graph showing the colocalization of calpastatin with NFH, measured using the Manders' coefficient (Kruskal-Wallis test).**(C)** Bar graph of the mean intensity ratio of calpastatin staining in the nucleus compared to the cytoplasm (Kruskal-Wallis test).

**Fig Supp 5. CAST compensation with CAST and Calp treatment improves survival in sporadic-ALS (sALS) MN cultures. (A)** Bar graph shows the relative expression of the endogenous calpastatin (*CAST*) gene. mRNA was normalized to *GAPDH* mRNA level (N=3, Kruskal-Wallis test). **(B)** Immunoblot of CAST protein level. Extractions were performed in MNs harvested after 3 weeks post-plating. **(C)** Representative images of MNs differentiated for 4 weeks stained for Calpastatin (green), Calpain2 (gray), and Neurofilament NF-H (red). Nuclei were counterstained with Hoechst (blue). Scale bar: 50µm. **(D)** Bar graphs of viability of MN cultures differentiated with and without NTFs at 4 weeks post-plating and after treatment with 5µM active CAST peptide or 10µM calpeptin for 24 hours. (N= 3, 1-way ANOVA). **(E-G)** Longitudinal changes in mean firing rate **(E)**, number of spikes by burst **(F)** and Burst Frequency average **(G)** of MNs recorded weekly over a span of 4 weeks post-plating. (N=1, Multiple t-test). NB: in the absence of burst, values have been artificially adjusted to zero. **(H-J)** Quantification de mean firing rate (**H**), number of spikes by burst (**I**) and Burst Frequency average (**J**) of 4 weeks post-plating MNs recorded 1 week after a 24hours-treatment (N=1, One-way-ANOVA). Data are represented as mean ± SEM. ns, not statistically significant; *$P < 0.05$, **$P < 0.01$, ***$P < 0.001$, ****$P<0.0001$. Data are represented as mean ± SEM. ns, not statistically significant; *$P < 0.05$.

**A**

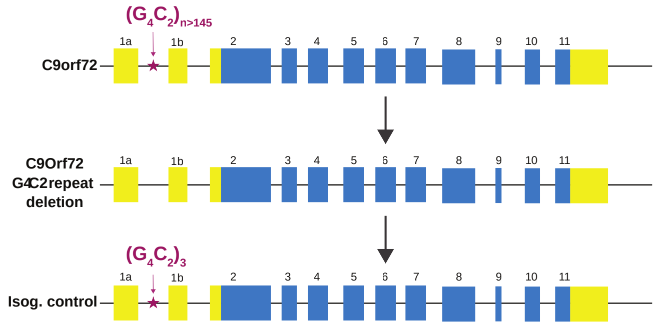


**B**

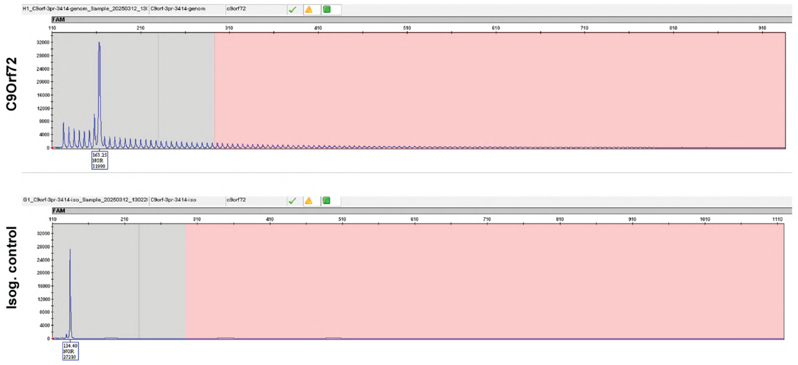


**C**

**Donor template**

**C9Orf72**
**G4C2repeat**
**deletion**

**Isog. control**

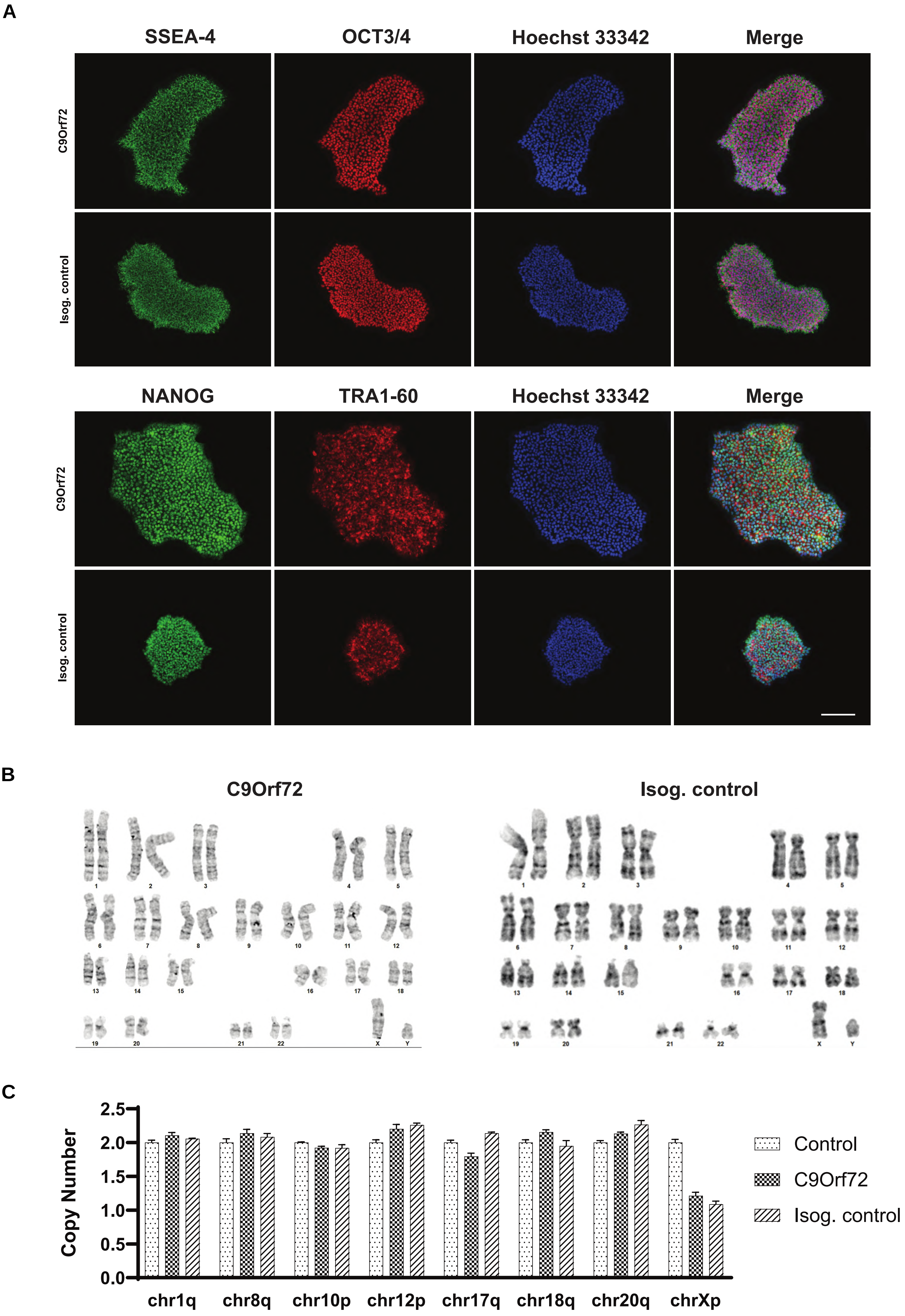

A
SSEA-4
OCT3/4
Hoechst 33342
Merge
C9Orf72
Isog. control
NANOG
TRA1-60
Hoechst 33342
Merge
C9Orf72
Isog. control
B
C9Orf72
Isog. control
C
Copy Number
2.5
2.0
1.5
1.0
0.5
0.0
chr1q
chr8q
chr10p
chr12p
chr17q
chr18q
chr20q
chrXp
Control
C9Orf72
Isog. control

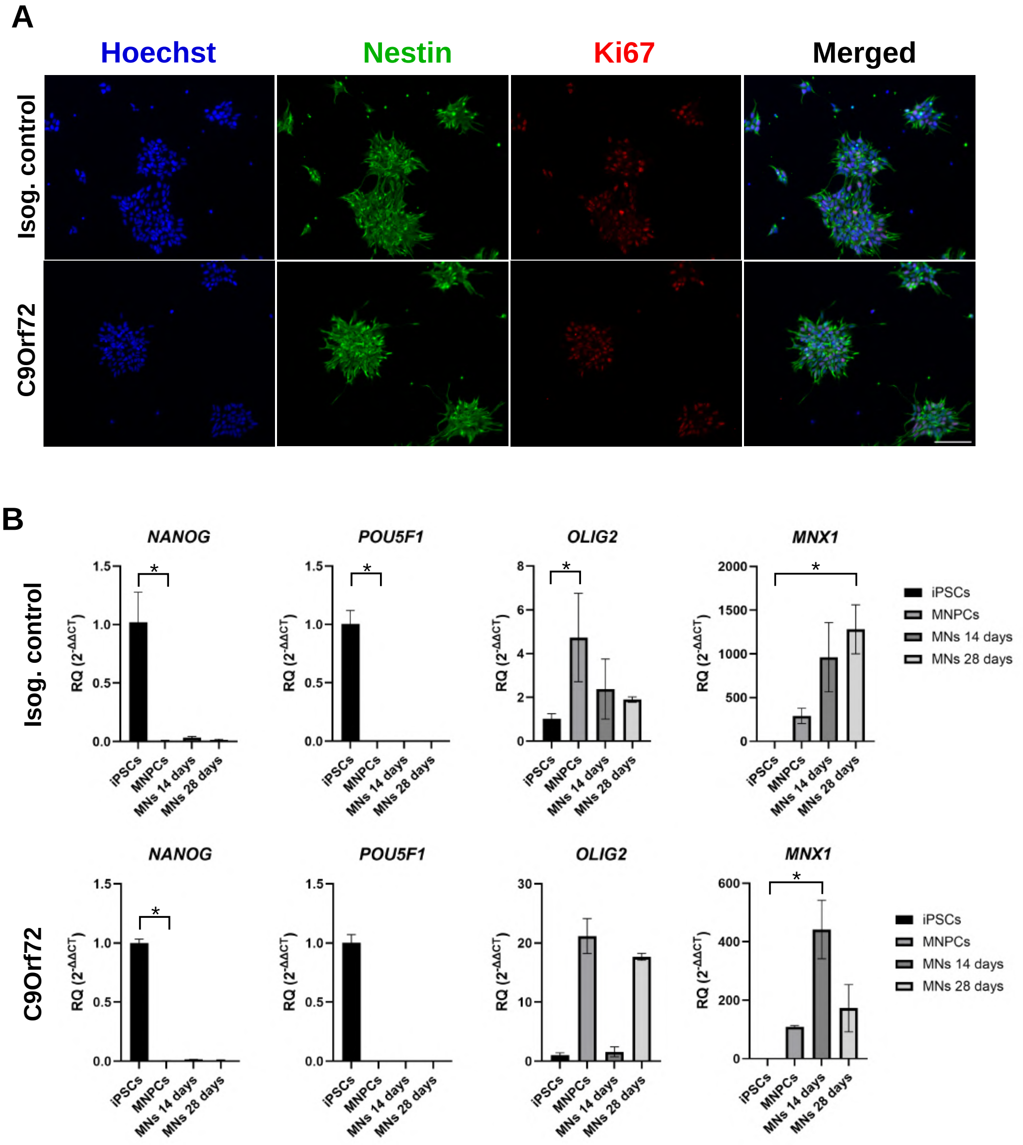
A
Hoechst
Nestin
Ki67
Merged
Isog. control
C9Orf72
B
NANOG
POU5F1
OLIG2
MNX1
RQ (2^-ΔΔCT)
iPSCs
MNPCs
MNs 14 days
MNs 28 days
*

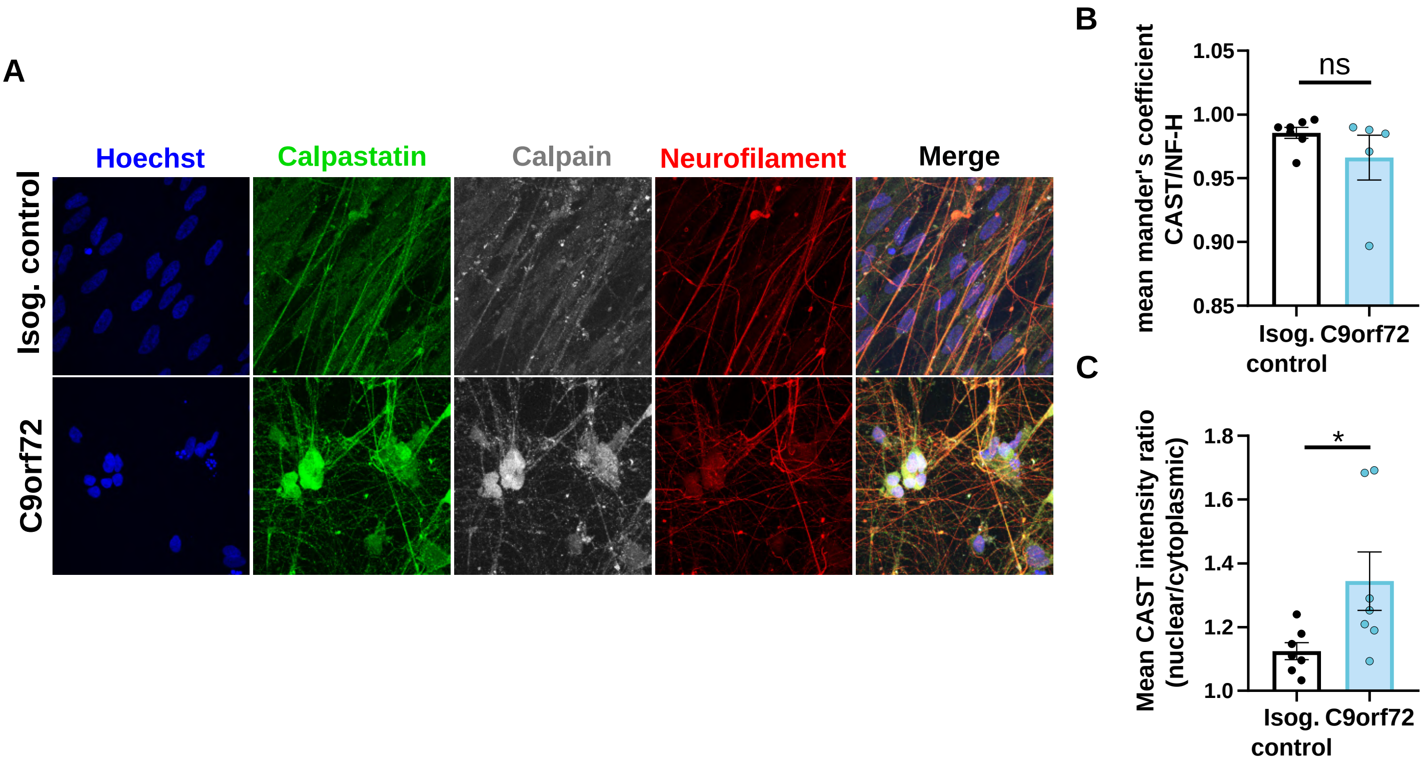
A
Hoechst
Calpastatin
Calpain
Neurofilament
Merge
Isog. control
C9orf72
B
mean mander's coefficient CAST/NF-H
ns
1.05
1.00
0.95
0.90
0.85
Isog. control
C9orf72
C
Mean CAST intensity ratio (nuclear/cytoplasmic)
*
1.8
1.6
1.4
1.2
1.0
Isog. control
C9orf72

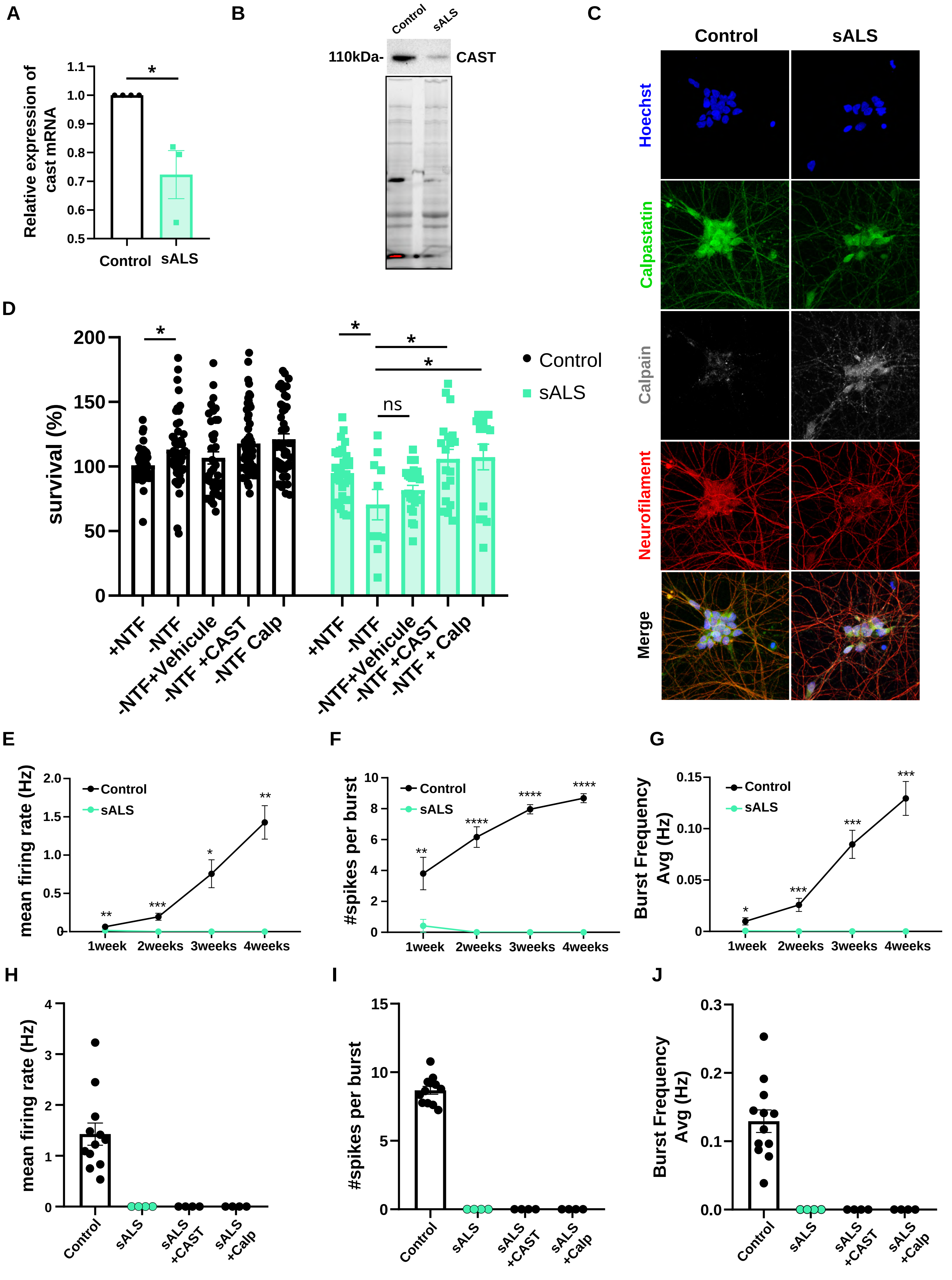

A
Relative expression of cast mRNA
Control
sALS
*
B
Control
sALS
110kDa-
CAST
C
Control
sALS
Hoechst
Calpastatin
Calpain
Neurofilament
Merge
D
survival (%)
Control
sALS
ns
+NTF
-NTF
-NTF+Vehicule
-NTF +CAST
-NTF Calp
-NTF + Calp
E
mean firing rate (Hz)
1week
2weeks
3weeks
4weeks
F
#spikes per burst
G
Burst Frequency Avg (Hz)
H
I
J
sALS +CAST
sALS +Calp
